# Plankton-infecting relatives of herpesviruses clarify the evolutionary trajectory of giant viruses

**DOI:** 10.1101/2021.12.27.474232

**Authors:** Morgan Gaïa, Lingjie Meng, Eric Pelletier, Patrick Forterre, Chiara Vanni, Antonio Fernandez-Guerra, Olivier Jaillon, Patrick Wincker, Hiroyuki Ogata, Mart Krupovic, Tom O. Delmont

**Author notes:** Co-first authors (Morgan Gaïa, Lingjie Meng).

## Abstract

DNA viruses have a major influence on the ecology and evolution of cellular organisms, but their overall diversity and evolutionary trajectories remain elusive. Here, we performed a phylogeny-guided genome-resolved metagenomic survey of the sunlit oceans and discovered plankton-infecting relatives of herpesviruses that form a putative new phylum dubbed ‘*Mirusviricota*’. The virion morphogenesis module of this large monophyletic clade is typical of viruses from the realm *Duplodnaviria*, with the major capsid protein fold being a likely structural intermediate between the capsid proteins of *Caudoviricetes* (tailed phages) and *Herpesvirales* (animal-infecting viruses). Yet, a substantial fraction of ‘*Mirusviricota’* genes, including hallmark transcription machinery genes missing in herpesviruses, are closely related homologs of large and giant eukaryotic DNA viruses from another viral realm. The remarkable chimeric attributes of ‘*Mirusviricota*’ provide missing links in the evolution of both herpesviruses and giant viruses. Furthermore, mirusviruses are widespread and transcriptionally active from pole to pole, encoding complex functional traits used during the infection of microbial eukaryotes. The ‘*Mirusviricota*’ prevalence, functional activity, diversification, and atypical evolutionary traits point to a lasting role of mirusviruses in the ecology of marine ecosystems that might have not only predated but also contributed to the emergence of herpesviruses and giant viruses.

## Introduction

Most double-stranded DNA viruses are classified into two major realms: *Duplodnaviria* and *Varidnaviria. Duplodnaviria* comprises tailed bacteriophages and related archaeal viruses of the class *Caudoviricetes* as well as eukaryotic viruses of the order *Herpesvirales. Varidnaviria* includes large and giant eukaryotic DNA viruses from the phylum *Nucleocytoviricota* as well as smaller viruses with tailless icosahedral capsid^1^. The two realms were established based on the non-homologous sets of virion morphogenesis genes (virion module), including the structurally unrelated major capsid proteins (MCPs) with the ‘double jelly-roll’ and HK97 folds in *Varidnaviria* and *Duplodnaviria*, respectively^1^. Both realms are represented across all domains of life, with the respective ancestors thought to date back to the last universal cellular ancestor^2^.

Within *Duplodnaviria*, bacterial and archaeal members of the *Caudoviricetes* display a continuous range of genome sizes, from ∼10 kb to >700 kb, whereas herpesviruses, restricted to animal hosts, are more uniform with genomes in the range of 100-300 kb. Herpesviruses likely evolved from bacteriophages, but the lack of related viruses outside of the animal kingdom raises questions regarding their exact evolutionary trajectory^3^. Members of the *Varidnaviria* also display a wide range of genome sizes, from ∼10 kb to >2 Mb, but there is a discontinuity in the complexity between large and giant viruses of the *Nucleocytoviricota* phylum and the rest of varidnaviruses with genomes <50 kb. It has been suggested that *Nucleocytoviricota* have evolved from a smaller varidnavirus ancestor^4–6^, but the complexification entailing acquisition of multiple informational genes (informational module) remains to be fully understood.

The *Caudoviricetes* and *Nucleocytoviricota* viruses are prevalent in the sunlit ocean where they play a critical role in regulating the community composition and blooming activity of plankton^7–13^. Here, we performed a genome-resolved metagenomic survey of planktonic DNA viruses guided by the phylogeny of a single hallmark gene. The survey covers nearly 300 billion metagenomic reads from surface ocean samples of the *Tara* Oceans expeditions^14–16^. We characterized and manually curated hundreds of *Nucleocytoviricota* population genomes that included a putative new class. But most notably, our survey led to the discovery of plankton-infecting relatives of herpesviruses that form a putative new phylum we dubbed ‘*Mirusviricota*’. The mirusviruses share complex functional traits and are widespread in the sunlit oceans where they actively infect eukaryotes, filling a critical gap in our ecological understanding of plankton. Furthermore, the remarkable chimeric attributes of ‘*Mirusviricota*’ provide key insights into the evolution of DNA viruses, clarifying the evolutionary trajectory of animal herpesviruses from tailed bacteriophages and giant viruses of the *Nucleocytoviricota* from smaller varidnaviruses.

## Results

### Environmental genomics of abundant marine eukaryotic DNA viruses

DNA–dependent RNA polymerase subunits A (RNApolA) and B (RNApolB) are evolutionarily informative gene markers occurring in most of the known DNA viruses infecting marine microbial eukartyotes^5,17^, which until now only included *Nucleocytoviricota*. Here, we performed a comprehensive search for RNApolB genes from the euphotic zone of polar, temperate, and tropical oceans using large co-assemblies from 798 metagenomes (total of 280 billion reads that produced ∼12 million contigs longer than 2,500 nucleotides)^15,16^ derived from the *Tara* Oceans expeditions^14^. These metagenomes encompass eight plankton size fractions ranging from 0.8 μm to 2000 μm (Table S1), all enriched in microbial eukaryotes^18,19^. We identified RNApolB genes in these contigs using a broad-spectrum hidden Markov model (HMM) profile and subsequently built a database of more than 2,500 non- redundant environmental RNApolB protein sequences (similarity <90%; Table S2). Phylogenetic signal for these sequences not only recapitulated the considerable diversity of marine *Nucleocytoviricota*^20^ but also revealed previously undescribed deep-branching lineages clearly disconnected from the three domains of life and other known viruses (Figure S1). We hypothesized that these new clades represent previously unknown lineages of double-stranded DNA viruses.

We performed a phylogeny-guided genome-resolved metagenomic survey focusing on the RNApolB of *Nucleocytoviricota* and new clades to manually delineate their genomic context (Table S3). We characterized and manually curated 587 non-redundant *Nucleocytoviricota* metagenome assembled genomes (MAGs) up to 1.45 Mbp in length (average of ∼270 Kbp) and 111 non-redundant MAGs up to 438 kbp in length (average of ∼200 Kbp) for the new clades. These MAGs were identified in all the oceanic regions (Table S3). We incorporated marine *Nucleocytoviricota* MAGs from previous metagenomic surveys^7,8^ and reference genomes from culture and cell sorting to construct a comprehensive database enriched in large and giant marine eukaryotic double-stranded DNA viruses (thereafter called Global Ocean Eukaryotic Viral [GOEV] database, Table S4). The GOEV database contains ∼0.6 million genes and provides contextual information to identify main ecological and evolutionary properties of MAGs containing to the new RNApolB clades.

### Mirusviruses are plankton-infecting relatives of herpesviruses

The newly assembled *Nucleocytoviricota* MAGs contain most of the hallmark genes of this viral phylum, corresponding to the virion and informational modules^4,5^ (Table S4). They expand the known diversity of the class *Megaviricetes* (grouping the *Imitervirales, Pandoravirales*, and *Algavirales* orders) and exposed a putative new class-level group we dubbed ‘*Proculviricetes*’ (represented by several MAGs exclusively detected in the Arctic and Southern Oceans) (Figure 1). MAGs from the new RNApolB clades also contain key genes evolutionarily related to the *Nucleocytoviricota* informational module, including RNApolA and RNApolB, family B DNA polymerase (DNApolB), and the transcription factor II-S (TFIIS; Figures 1 and S2). For instance, the RNApolA of most of these MAGs branched as a sister clade to the order ‘*Pandoravirales’*. Phylogenomic inferences of the concatenated four informational gene markers (see Figure 1) indicate that they represent a new viral clade with several hallmark genes closely related to, yet distinct from those in the known *Nucleocytoviricota* classes. We dubbed viruses in this monophyletic clade the mirusviruses (Mirus is a Latin word for surprising, strange). Their MAGs are organized into seven distinct subclades, M1 to M7 (from the most to least populated), with the latter being represented by a single MAG (Figure 2A, Table S4). Notably, however, the mirusvirus MAGs were devoid of identifiable homologs of the *Nucleocytoviricota* virion module, including the double jelly-roll MCP (Figure 1). Instead, annotation of mirusvirus gene clusters using sensitive sequence and structure similarity searches (see Methods) identified a distant homolog of HK97-fold MCPs occurring in most of these MAGs (Figure 1). This MCP fold, only shared with *Caudoviricetes* and *Herpesvirales*, indicates that mirusviruses belong to the realm *Duplodnaviria*.

**Figure 1:**
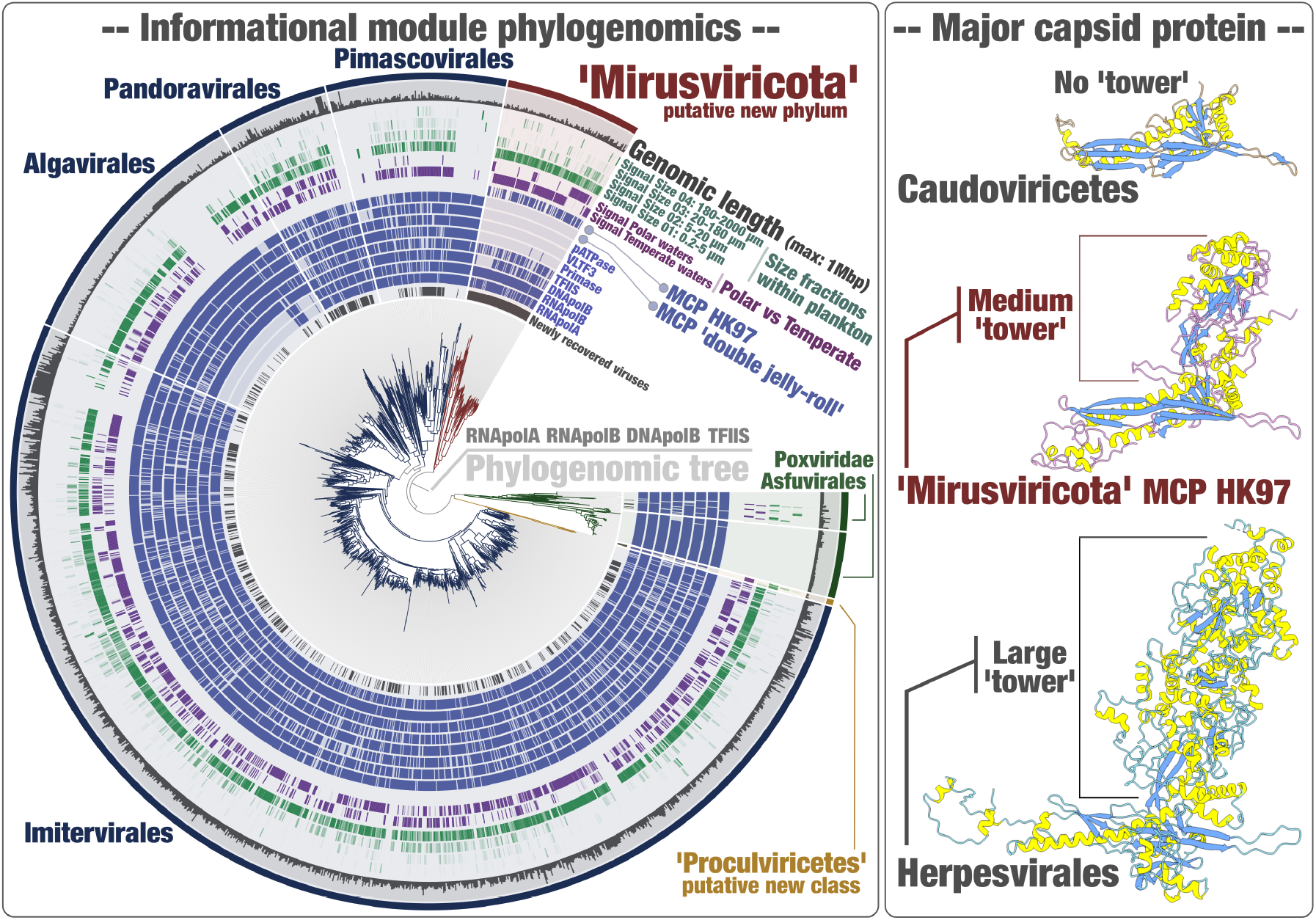
Evolutionary relationships between *Nucleocytoviricota, Herpesvirales* and mirusviruses. Left panel displays a maximum-likelihood phylogenetic tree built from the GOEV database (1,722 genomes) based on a concatenation of manually curated RNApolA, RNApolB, DNApolB and TFIIS genes (3,715 amino acid positions) using the PMSF mixture model and rooted between the *Pokkesviricetes* and the rest. The tree was decorated with layers of complementary information and visualized with anvi’o. Right panel displays MCP 3D structures for *Escherichia* phage HK97 (*Caudoviricetes*), a reference genome for the mirusviruses (obtained using AlphaFold2), and the human cytomegalovirus (*Herpesvirales*).

**Figure 2:**
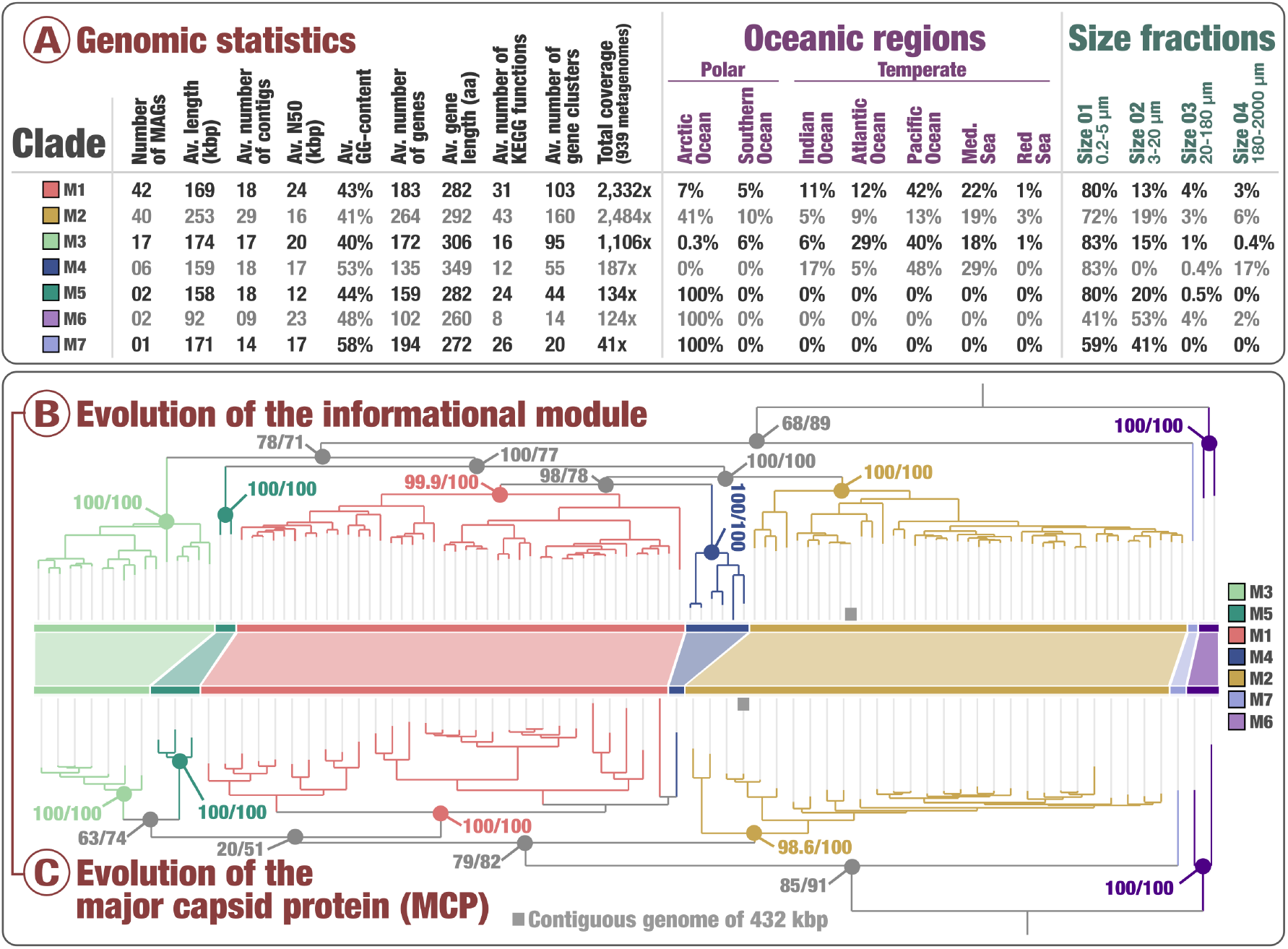
Genomic statistics and evolution of mirusviruses. Panel A displays genomic and environmental statistics for the seven ‘*Mirusviricota*’ clades. Panel B displays a maximum-likelihood phylogenetic tree built from the ‘*Mirusviricota*’ MAGs based on a concatenation of four hallmark informational genes (RNApolA, RNApolB, DNApolB, TFIIS; 3,715 amino acid positions) using the LG+F+R7 model. Panel C displays a maximum-likelihood phylogenetic tree built from the ‘*Mirusviricota*’ MAGs based on the Major Capsid Protein (701 amino acid positions) using the LG+R6 model. Both trees were rooted between clade M6 and other clades. Values at nodes represent branch supports (out of 100) calculated by the SH-like aLTR (1,000 replicates; left score) and ultrafast bootstrap approximations (1,000 replicates; right score).

In herpesvirus MCPs, the HK97-fold domain, referred to as the ‘floor’ domain and responsible for capsid shell formation, is embellished with a ‘tower’ domain that projects away from the surface of the assembled capsid^21^. The ‘tower’ domain is an insertion within the A-subdomain of the core HK97-fold^21,22^. In mirusviruses, the MCP protein also contains an insertion within the A-subdomain. Such ‘tower’ domain has not been thus far described for any member of the *Caudoviricetes*, including the so-called jumbo phages (i.e., phages with very large genome^23^). Considering the size and position of the mirusvirus MCP ‘tower’, it is possible that this protein is an evolutionary intermediate between *Caudoviricetes* (no tower) and *Herpesvirales* (larger tower) MCPs (Figures 1 and S3). However, the extent of protein sequence divergences prevents meaningful phylogenetic analyses to unequivocally determine the directionality of the MCP evolution. Yet, phylogenetic inferences of the mirusvirus MCP alone recapitulated the seven subclades initially identified based on the informational module (Figure 2B), indicative of a coevolution of the two functional modules. Finally, consistent with the identification of the HK97-fold MCP, further comparisons of the profile HMMs and predicted 3D structures uncovered the remaining key components of the *Duplodnaviria*-specific virion module, including the terminase (ATPase-nuclease, key component of the DNA packaging machine), portal protein, capsid maturation protease as well as triplex capsid proteins 1 and 2. The presence of these genes in mirusviruses establishes they are *bona fide* large DNA viruses capable of forming viral particles similar to those of previously known viruses in the realm *Duplodnaviria*.

Phylogenetic inferences of the DNApolB gene using the GOEV database and a wide range of eukaryotic and additional viral lineages^24^ supported the evolutionary distance of mirusviruses relative to all other known clades of double-stranded DNA viruses (Figure S4). The mirusvirus DNApolB was positioned at the base of a clade grouping the *Herpesviridae* and the eukaryotic Zeta-type sequences, and close to the eukaryotic Delta-type sequences and the class *Megaviricetes* of *Nucleocytoviricota* (with the closest relative being ‘*Pandoravirales*’). By contrast, they are distant from both *Caudoviricetes* and the class *Pokkesviricetes* of *Nucleocytoviricota*. Taken together, the (i) considerable genetic distances between the virion modules of mirusviruses, *Caudoviricetes* and *Herpesvirales* (preventing robust and informative phylogenetic inferences), (ii) distinct 3D structure of the mirusvirus MCP, (iii) and DNApolB phylogenetic inferences, firmly characterize mirusviruses as a putative new phylum, which we dubbed ‘*Mirusviricota*’. These viruses would hence represent a third clade within the realm *Duplodnaviria*, next to the well-known *Caudoviricetes* (phylum *Uroviricota*) and *Herpesvirales* (phylum *Peploviricota*) lineages.

### Mirusviruses have a unique and complex functional lifestyle

The 111 ‘*Mirusviricota’* MAGs contain a total of 22,242 genes organized into 35 core gene clusters present in at least 50% of MAGs, 1,825 non-core gene clusters, and finally 9,018 singletons with no close relatives within the GOEV database (Tables S5). Core gene clusters provided a window into critical functional capabilities shared across subclades of mirusviruses. Aside from the capsid and informational modules, they correspond to functions related to DNA stability (H3 histone), DNA replication (DNA replication licensing factor, glutaredoxin/ribonucleotide reductase, Holliday junction resolvase, three-prime repair exonuclease 1), transcription (multisubunit RNA polymerase, TATA-binding protein), gene expression regulation (lysine specific histone demethylase 1A), post-transcriptional modification of RNA (RtcB-like RNA-splicing ligase) and proteins (putative ubiquitin protein ligase), protein degradation (trypsin-like, C1 and M16-family peptidases), cell growth control (Ras-related protein), detection of external signals (sensor histidine kinase) and light sensitive receptor proteins (heliorhodopsins). Thus, mirusviruses encode an elaborate toolkit which could enable fine-tuning the cell biology and energetic potential of their hosts for optimal virus replication. Finally, ten core gene clusters could not be assigned any function, representing as many hallmark ‘*Mirusviricota’* genes coding for proteins (with confident 3D structure predictions distant from those in reference databases) that await experimental functional characterizations.

Clustering of ‘*Mirusviricota’* MAGs and reference viral genomes from culture (including *Nucleocytoviricota, Herpesvirales* and *Caudoviricetes*) based on quantitative occurrence of gene clusters highlighted the strong functional differentiation between mirusviruses and herpesviruses and, conversely, a strong functional similarity between mirusviruses and the class *Megaviricetes* of *Nucleocytoviricota* that is also widespread at the surface of the oceans (Figure S5 and Table S6). Thus, function-wise and aside from the virion morphogenesis module, mirusviruses more closely resemble the marine *Nucleocytoviricota* viruses as compared to *Herpesvirales*. To further explore the functional landscape of eukaryote-infecting marine viruses, we clustered their genomes based on quantitative occurrence of gene clusters using the entire GOEV database (Table S5). The mirusviruses clustered together and were further organized into subclades in line with phylogenomic signals (Figure S6). In contrast, this analysis emphasized the complex functional makeup of *Nucleocytoviricota* lineages, with some clades (e.g., the *Imitervirales* and *Algavirales*) split into multiple groups. Aside from the informational module, gene clusters connecting a substantial portion of ‘*Mirusviricota*’ and *Nucleocytoviricota* genomes were dominated by functions involved in DNA replication: the glutaredoxin/ribonucleotide reductase, Holliday junction resolvase, proliferating cell nuclear antigen, dUTPase and DNA topoisomerase II. Commonly shared functions also included the Ras protein, patatin-like phospholipase (lipid degradation), peptidase C1, ubiquitin carboxyl-terminal hydrolase (protein activity regulation) and Evr1–Alr family (maturation of cytosolic Fe/S protein). Thus, the functional connectivity between the two phyla goes well beyond the informational module. On the other hand, hundreds of gene clusters and functions were significantly enriched in either mirusviruses or *Nucleocytoviricota* (Tables S5 and S7), exposing distinct lifestyles for the two clades. Aside from the virion morphogenesis module, core gene clusters among the mirusviruses that were significantly less represented among *Nucleocytoviricota* genomes included the trypsin-like (73% of genomes in mirusviruses *vs*. 9% in *Nucleocytoviricota*) and M16-family (60% *vs*. 2%) peptidases, TATA-binding protein (59% *vs*. 0%), heliorhodopsin (64% *vs*. 5%) and histone (54% *vs* 2%). Notably, phylogenetic inferences of the histones and rhodopsins showed several monophyletic clades for ‘*Mirusviricota*’ relatively distant from the corresponding homologs in both *Nucleocytoviricota* and marine planktonic eukaryotes (Figure 3). Thus, these important genes are specific to mirusviruses and likely have been acquired a long time ago.

**Figure 3:**
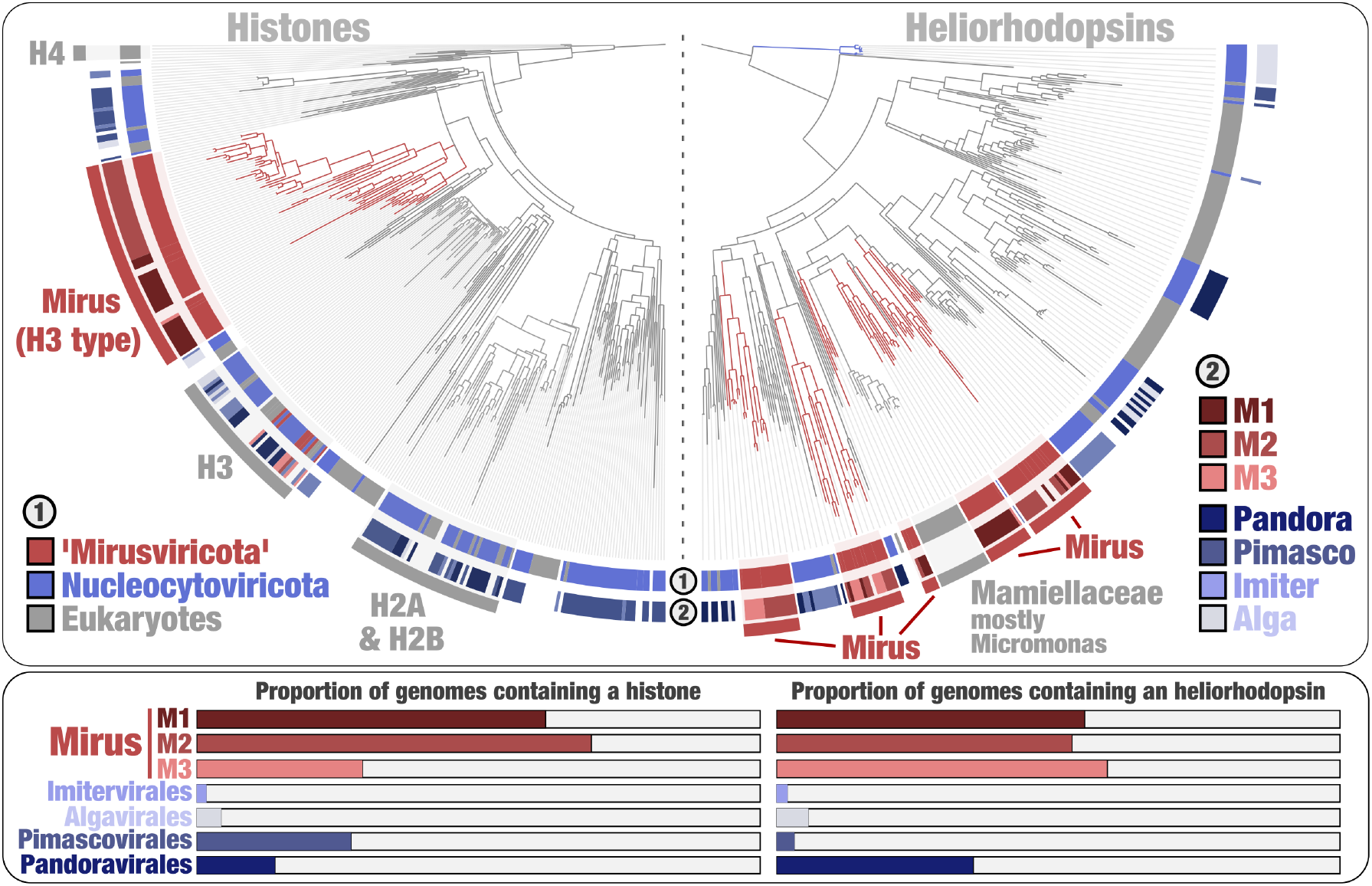
Mirusviruses contain new phylogenetic clades of histones and heliorhodopsins. Left panel displays a maximum-likelihood phylogenetic of histones occurring in the GOEV database and in eukaryotic MAGs, rooted with H4. The various eukaryotic clades distant from H2-H3-H4 were excluded to focus on the more restrained viral signal. Right panel displays a maximum-likelihood phylogenetic tree of heliorhodopsins occurring in the GOEV database and in eukaryotic MAGs, rooted with Algavirales clade. Bottom panels summarize the proportion (from 0 to 100%) of genomes from different viral clades containing histones and rhodopsins.

### Mirusviruses are widespread and actively infect microbial eukaryotes

To our knowledge, ‘*Mirusviricota*’ represents the first eukaryote-infecting lineage of *Duplodnaviria* found to be widespread and abundant within plankton in the sunlit ocean. Indeed, mirusviruses were detected in 131 out of the 143 *TARA* Oceans stations. They occurred mostly in the 0.2-5 μm (76.3% of the entire mirusvirus metagenomic signal) and 3-20 μm (15.4%) size fractions that cover a high diversity of unicellular planktonic eukaryotes^18^ (Figures 1 and 2, Table S8). As a first insight into virus-host interactions, phylogeny-guided host predictions^18^ based on co-occurrence patterns using ‘*Mirusviricota*’ and eukaryotic MAGs characterized from the same samples^13^ showed significant clade-level associations between subclade M1 and two orders of phytoplankton: Chloropicales (green algae) and Phaeocystales (haptophytes; Figure S7 and Table S8). In addition, heliorhodopsins encoded by *Micromonas* populations (prominent green algae widespread in the sunlit ocean) might have been acquired from mirusviruses through an ancient horizontal gene transfer (Figure 3). For now, precise host assignment for ‘*Mirusviricota*’ is lagging far behind genomic recoveries, echoing the knowledge gap for most *Nucleocytoviricota* viruses. Nonetheless, the two lines of evidence presented above suggest that mirusviruses have a long history of infecting abundant phytoplankton lineages.

The ‘*Mirusviricota*’ genes recruited 6.7 million metatranscriptomic reads from *Tara* Oceans (0.0016% of 420 billion reads), representing 13% of the overall signal for the GOEV database (Table S9). Thus, *in situ* transcriptomic signal for the 111 mirusvirus MAGs is already non-negligible as compared to the more numerous *Nucleocytoviricota* genomes (1,269 of them were detected in the *Tara* Oceans metagenomes), stressing the relevance of ‘*Mirusviricota’* to eukaryotic virus-host dynamics in marine systems. Mirusviruses were most active in the sunlit ocean (and especially in the euphotic subsurface layer enriched in chlorophyll) as compared to the mesopelagic zone (> 200m in depth), and within the cellular range of 0.2-20 μm (Figure 4), in line with the metagenomic signal. The 35 core gene clusters for ‘*Mirusviricota’* represented 20% of the metatranscriptomic signal (including 12% for just seven capsid proteins), with remaining signal linked to non-core gene clusters (43%) and singletons (37%). Thus, highly diversified genes (nearly 10,000 singletons were identified) appear to play a critical role in the functional activity of ‘*Mirusviricota’* during infection of marine microbial eukaryotes. Some of those genes might contribute to countering the defense mechanisms as part of a longstanding co-evolution with the hosts.

**Figure 4:**
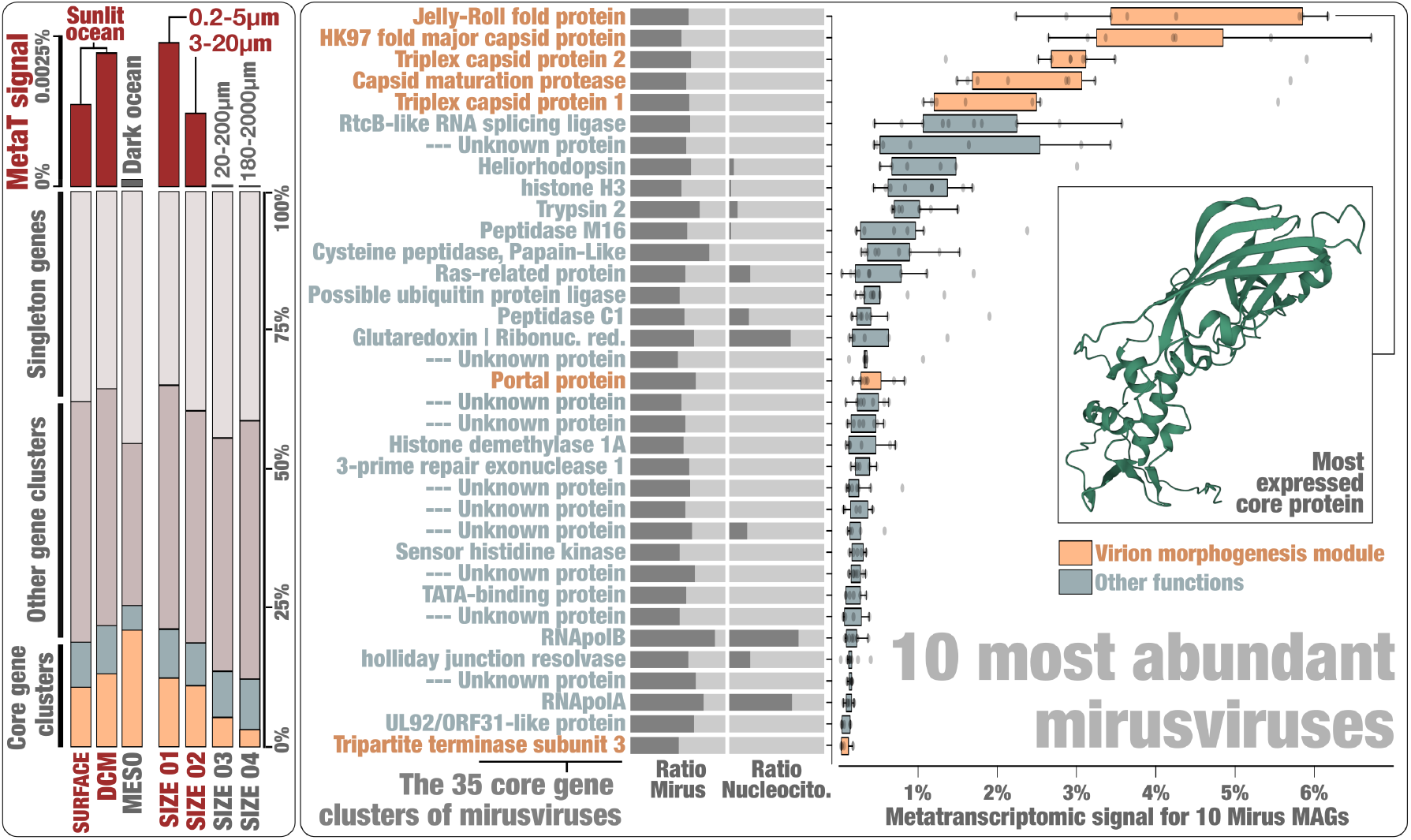
*In situ* expression profile of mirusviruses during infection. Left panel summarizes the overall metatranscriptomic signal for all 111 Mirus MAGs among different categories of metatranscriptomes (e.g., surface samples versus). DCM and MESO stand for Deep Chlorophyll maximum layer and Mesopelagic (top dark ocean layer below 200 meters). In addition, bars show the cumulated mean coverage signal of 3 gene categories (the 35 core gene clusters, non-core gene clusters and singletons). Core gene clusters were further split into those related to the virion morphogenesis module in orange and other functions (including unknown core gene clusters) in blue. Right panel summarizes the occurrence of 35 ‘*Mirusviricota*’ core gene clusters among the Mirus MAGs and *Nucleocytoviricota* genomes (ratio of genomes containing the gene cluster, from 0 to 1). The panel then displays boxplots corresponding to the overall metatranscriptomic signal for genes corresponding to the core gene clusters and occurring in the ten most abundant mirusviruses within the scope of *Tara* Oceans metagenomes. Percentage values are genome-centric and correspond to the percentage of mean coverage (sum across all the metatranscriptomes) of one gene when considering to the cumulated mean coverage of all genes (sum across all the metatranscriptomes) found in the corresponding genome.

Mirusviruses have different ecological niches (e.g., some are only found in the Arctic Ocean), yet their 35 core genes were expressed with similar levels in samples with metatranscriptomic signal, indicating a relatively homogeneous functional lifestyle regardless of latitude or subclade (Figure 4 and Table S9). Genes coding for the capsid proteins were the most expressed, with ratios recapitulating the proportion of corresponding proteins in the capsid of herpesviruses (e.g., more HK97 MCPs as compared to triplex or portal proteins). Genes coding for the new types of heliorhodopsin and histone were also substantially expressed, pointing to an important functional role during infection. In comparison, the RNApolA and RNApolB systematically displayed lower levels of expression during sampling by the *Tara* Oceans expeditions. Collectively, the biogeographic and *in situ* transcriptomic patterns of mirusviruses suggest they actively infect abundant marine unicellular eukaryotes in both temperate and polar waters.

### The mirusvirus informational module links two major realms of DNA viruses

To further validate the genomic content of mirusviruses and to exclude the possibility of artificial chimerism, we created an HMM for the newly identified ‘*Mirusviricota’* MCP and used it as bait to search for complete genomes in additional databases. First, we only found two ‘*Mirusviricota’* MCPs in a comprehensive viral genomic resource from the <0.2 μm size fraction of the surface oceans (GOV2^12^), suggesting that most virions in this clade are larger than 0.2 μm in size. We subsequently screened for the ‘*Mirusviricota’* MCP in a database containing hundreds of metagenomic assemblies from the 0.2-3 μm size fraction of the surface oceans^25^. We found a contiguous ‘*Mirusviricota*’ genome (355 genes) in the Mediterranean Sea affiliated to the clade M2 with a length of 431.5 kb, just 6 kb shorter than the longest ‘*Mirusviricota’* MAG (Figure 2, panels B and C). Its genes recapitulate the core functionalities of mirusviruses (e.g., topoisomerase II, TATA-binding protein, histone, multiple heliorhodopsins, Ras-related GTPases, cell surface receptor, ubiquitin, and trypsin), and 80 of these genes have a clear hit when compared to *Nucleocytoviricota* HMMs (see Figure S8 and Methods). This genome also contains a 34.6 kb-long non-core gene of unknown function that might represent a record holder in terms of length for a viral gene. Most critically, all hallmark genes for the informational (DNApolB, RNApolA, RNApolB, TFIIS) and virion (MCP HK97, terminase, portal protein, capsid maturation protease, the two triplex capsid proteins, and jelly-roll fold protein) modules of ‘*Mirusviricota’* are not only present but also occur relatively homogeneously across the genome (Figure S8). Thus, this near-complete contiguous genome perfectly recapitulated all apparent chimeric attributes of ‘*Mirusviricota*’.

On the one hand, mirusviruses belong to the realm *Duplodnaviria* based on their virion module. On the other hand, we also showed that hallmark informational genes (some of which are missing in herpesviruses) display surprisingly high sequence similarity to the corresponding genes prevalent in the phylum *Nucleocytoviricota*. These results strongly indicate that this informational module was once transferred from one realm to the other, most likely after the long-standing co-evolution of the corresponding genes between viruses and proto-eukaryotic hosts^5^ (Figure 5). Importantly, the *Nucleocytoviricota* viruses (including all known giant viruses) would be chimeric in the case of a *Duplodnaviria* origin of the informational module. Besides, our data indicates that herpesviruses could have emerged from a mirusvirus ancestor through reductive evolution that involved loss of the transcription machinery. Thus, the mirusviruses are not only integral components of the ecology of eukaryotic plankton, but they also fill critical gaps in our understanding of the evolutionary trajectories of two major realms of double-stranded DNA viruses.

**Figure 5:**
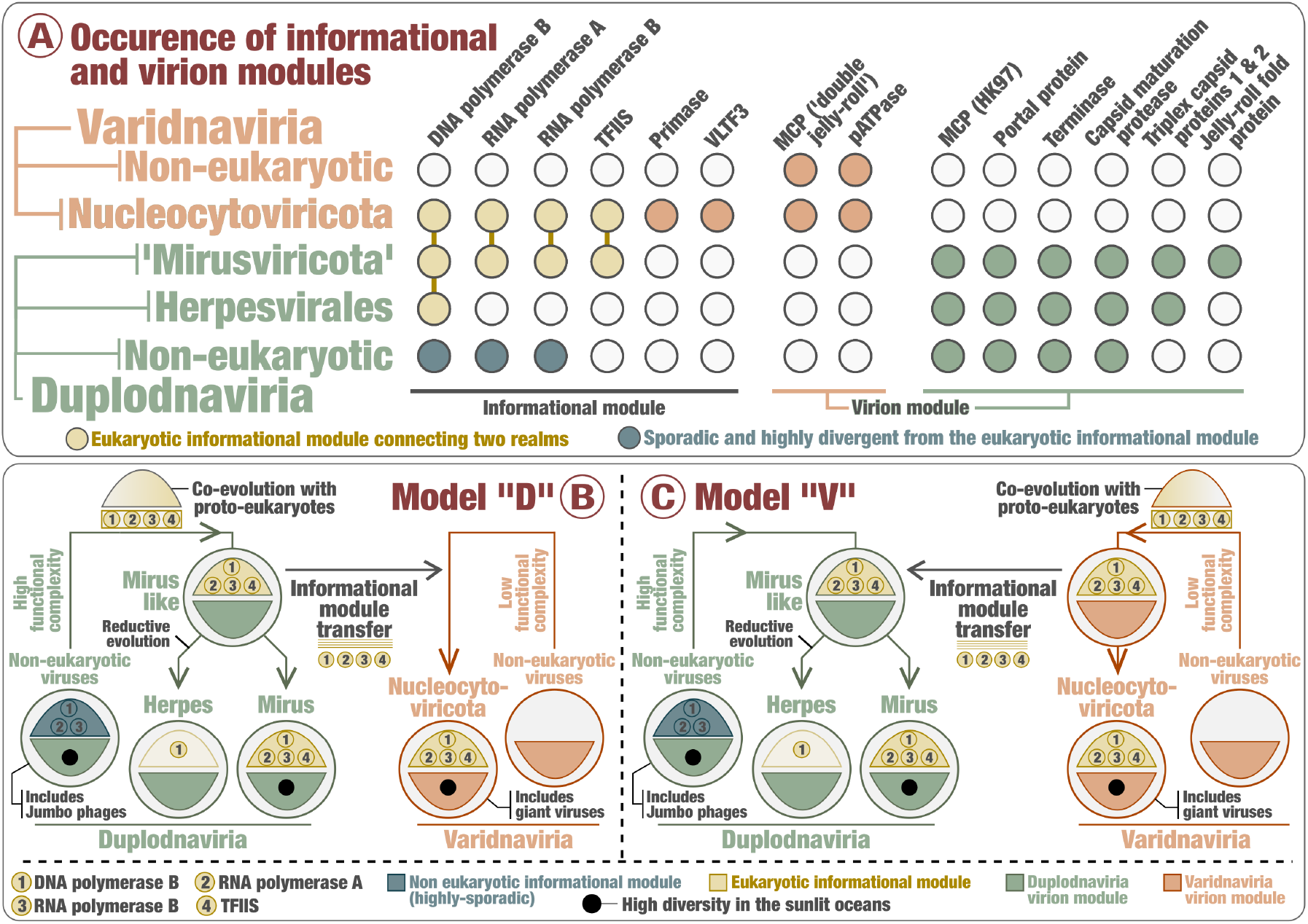
Evolutionary trajectories of the eukaryotic informational module. Panel A summarizes the occurrence of hallmark genes for the informational and virion modules in *Nucleocytoviricota*, mirusviruses, herpesviruses and non-eukaryotic genomes from the two realms. Panels B and C describe two evolutionary scenarii in which the informational module of eukaryotic infecting viruses within the realms *Duplodnaviria* and *Varidnaviria* first emerged in the ancestor of either mirusviruses (Model “D”) or *Nucleocytoviricota* (Model “V”).

## Discussion

Our phylogeny-guided genome-resolved metagenomic survey of plankton in the surface of five oceans and two seas exposed a major clade of large eukaryotic DNA viruses, with genomes up to >400 Kbp in length, that are diverse, prevalent, and active in the sunlit ocean. This clade, dubbed ‘*Mirusviricota’*, corresponds to a putative new phylum within the realm *Duplodnaviria* that until now only included the bacteria- and archaea-infecting *Caudoviricetes* and animal-infecting *Herpesvirales*. The ‘*Mirusviricota*’ phylum is organized into at least seven subclades that might correspond to distinct families. Although both mirusviruses and *Herpesvirales* are eukaryote-infecting duplodnaviruses, they display very different genomic features. Most notably, mirusviruses substantially deviate from all other previously characterized groups of DNA viruses, with the virion morphogenesis module (a basis for highest-rank double-stranded DNA virus taxonomy) affiliated to the realm *Duplodnaviria* and the informational module closely related to that of large and giant viruses within the realm *Varidnaviria*. These apparent chimeric attributes were recapitulated in a near-complete contiguous genome of 431.5 Kbp. ‘*Mirusviricota*’ has a substantial functional overlap with large and giant eukaryotic varidnaviruses that goes well beyond the informational module and includes ecosystem- and host-specific genes, which could have been horizontally transferred between the two groups of viruses or convergently acquired from the shared hosts. The discovery of ‘*Mirusviricota*’ is a reminder that we have not yet grasped the full ecological and evolutionary complexity of even the most abundant double-stranded DNA viruses in key ecosystems such as the surface of our oceans and seas.

Mirusviruses are relatively abundant in various regions of the sunlit oceans where they actively infect eukaryotic plankton smaller than 20 μm in size and express a variety of functions. Viral-host predictions linked some mirusviruses to several eukaryotic lineages including key phototrophs (e.g., haptophytes) that now await experimental validations. In terms of functions, mirusviruses have a cohesive and rather complex inferred lifestyle (wide array of functions) that resembles, to some extent, that of marine varidnaviruses^7,8^. For instance, the patatin-like phospholipase shared between the two phyla had already been suggested to promote the transport of *Nucleocytoviricota* genomes to the cytoplasm and nucleus^26^. Functions enriched in mirusviruses as compared to the *Nucleocytoviricota* include phylogenetically distinct H3 histones (proteins involved in chromatin formation within the eukaryotic cells^27^) and heliorhodopsins (light sensitive receptor proteins that can be used as proton channels by giant viruses during infection^28^). Interestingly, a core *Micromonas* heliorhodopsin may have originated from a mirusvirus, implicating them as important players in the evolution of planktonic eukaryotes by means of gene flow. Concurrently, phylogenetic relationship between mirusvirus and *Micromonas* heliorhodopsins suggests that green algae might have served as hosts for some of the ‘*Mirusviricota*’ lineages. Together, biogeographic patterns, functional gene repertoires, and metatranscriptomic signal indicate that mirusviruses influence the ecology of key marine eukaryotes using a previously overlooked lifestyle.

Viruses of the *Herpesvirales* and *Nucleocytoviricota* belong to two ancient virus lineages, *Duplodnaviria* and *Varidnaviria*, respectively, with their corresponding ancestors possibly antedating the last universal cellular ancestor (LUCA)^1,2^. Nevertheless, the exact evolutionary trajectories and the identity of the respective most recent common ancestors of these prominent eukaryote-infecting double-stranded DNA viral clades remained elusive, in part due to the lack of known intermediate states. Particularly puzzling is the evolutionary trench between relatively simple varidnaviruses (those infecting Bacteria and Archaea, as well as virophages and *Adenoviridae*), with modest gene repertoires for virion formation and genome replication, and complex eukaryotic varidnaviruses (the *Nucleocytoviricota*) with large and giant genomes encoding multiple functions, including nearly complete genome replication and transcription machinery. Similarly enigmatic is the gap between the ubiquitous *Caudoviricetes*, some of which rival *Nucleocytoviricota* in terms of functional complexity and richness of their gene repertoires, and *Herpesvirales*, which are restricted to animal hosts and uniformly lack the transcription machinery and practice nuclear replication. The apparent chimeric attributes of ‘*Mirusviricota’* might now help to clarify both of these long-standing questions. It is possible that genome complexity observed in the contemporary ‘*Mirusviricota*’ lineages has been attained and inherited from a common ancestor with large *Caudoviricetes*. Indeed, most of the proteins involved in genome replication and nucleotide metabolism conserved in mirusviruses and *Nucleocytoviricota* are also sporadically occurring in *Caudoviricetes* (e.g., divergent RNApolA and RNApolB genes^29^). It is possible that these proteins were later transferred between the ancestors of ‘*Mirusviricota’* and *Nucleocytoviricota*, a scenario consistent with a closer sequence similarity between the two virus groups. The transfer of the informational module from a mirusvirus to the ancestor of *Nucleocytoviricota* would explain a sudden transition from ‘small’ varidnaviruses to complex *Nucleocytoviricota*. However, a transfer from *Nucleocytoviricota* to the ancestor of mirusviruses cannot be excluded at this stage. Beside the core functions responsible for virion formation and information processing, another important component of both *Nucleocytoviricota* and ‘*Mirusviricota*’ genomes, specifically related to eukaryote-specific aspects of virus-host interactions, has likely evolved more recently through horizontal gene transfers or convergent gene acquisition from different sources, facilitated by shared hosts and ecosystems. Finally, the identification of ‘*Mirusviricota*’ extends the presence of duplodnaviruses beyond animals to eukaryotic plankton hosts, strongly suggesting their ancient association with eukaryotes. The presence and location of the tower domain combined with the conservation of the two triplex capsid proteins (none of these are present in known *Caudoviricetes*) in both ‘*Mirusviricota*’ and *Herpesvirales* suggest a common ancestry, rather than independent evolution from distinct *Caudoviricetes* clades. With the shorter size of the tower domain and considering later emergence of animals compared to unicellular eukaryotes, ‘*Mirusviricota*’ viruses might more closely resemble the ancestral state of eukaryotic duplodnaviruses. If so, animal herpesviruses would have evolved by reductive evolution^1,2^.

Overall, the prevalence, functional complexity (including novel core proteins of unknown function) and verified transcriptional activity of ‘*Mirusviricota*’ point to a lasting role of the mirusviruses in the ecology of marine ecosystems that might have not only predated, but also contributed to the emergence of herpesviruses and large and giant viruses from another realm. Thus, this putative phylum has expanded our understanding of plankton ecology and viral evolution. Moving forward, additional functional and genomic characterizations coupled with cell sorting for host predictions will contribute to further assessing the lifestyle and prominence of mirusviruses within the oceans and beyond.

## Material & methods

### *Tara* Oceans metagenomes and metatranscriptomes

We analyzed 937 metagenomes and 1,149 metatranscriptomes from *Tara* Oceans available at the EBI under project PRJEB402 (https://www.ebi.ac.uk/ena/browser/view/PRJEB402). Tables S1 and S9 report general information (including the number of reads and environmental metadata) for each metagenome and metatranscriptome.

### Constrained automatic binning with CONCOCT

The 798 metagenomes corresponding to size fractions ranging from 0.8 μm to 2 mm were previously organized into 11 ‘metagenomic sets’ based upon their geographic coordinates^15,16^. Those 0.28 trillion reads were used as inputs for 11 metagenomic co-assemblies using MEGAHIT^30^ v1.1.1, and the contig header names were simplified in the resulting assembly outputs using anvi’o^31^ v6.1. Co-assemblies yielded 78 million contigs longer than 1,000 nucleotides for a total volume of 150.7 Gbp. Constrained automatic binning was performed on each co-assembly output, focusing only on the 11.9 million contigs longer than 2,500 nucleotides. Briefly, (1) anvi’o profiled contigs using Prodigal^32^ v2.6.3 with default parameters to identify an initial set of genes, (2) we mapped short reads from the metagenomic set to the contig using BWA v0.7.15^33^ (minimum identity of 95%) and stored the recruited reads as BAM files using samtools^34^, (3) anvi’o profiled each BAM file to estimate the coverage and detection statistics of each contig, and combined mapping profiles into a merged profile database for each metagenomic set. We then clustered contigs with the automatic binning algorithm CONCOCT^35^ by constraining the number of clusters per metagenomic set to a number ranging from 50 to 400 depending on the set (total of 2,550 metagenomic blocks from ∼12 million contigs).

### Diversity of DNA-dependent RNA polymerase B subunit genes

We used HMMER ^36^ v3.1b2 to detect genes matching to the DNA-dependent RNA polymerase B subunit (RNApolB) among all 2,550 metagenomic blocks based on a single HMM model. We used CD-HIT^37^ to create a non-redundant database of RNApolB genes at the amino acid level with sequence similarity <90% (longest hit was selected for each cluster). Short sequences were excluded. Finally, we included reference RNApolB amino acid sequences from Bacteria, Archaea, Eukarya and giant viruses^5^: the sequences were aligned with MAFFT^38^ v7.464 and the FFT-NS-i algorithm with default parameters and trimmed at >50% gaps with Goalign v0.3.5 (https://www.github.com/evolbioinfo/goalign). We performed a phylogenetic reconstruction using the best fitting model according to the Bayesian Information Criterion (BIC) from the ModelFinder^39^ Plus option with IQ-TREE^40^ v1.6.2. We visualized and rooted the phylogeny using anvi’o. This tree allowed us to identify new RNApolB clades.

### Phylogeny-guided genome-resolved metagenomics

Each metagenomic block containing at least one of the RNApolB genes of interest (see previous section) was manually binned using the anvi’o interactive interface to specifically search for *Nucleocytoviricota* and Mirus MAGs. First, we used HMMER^36^ v3.1b2 to identify eight hallmark genes (eight distinct HMM runs within anvi’o) as well 149 additional orthologous groups often found in reference *Nucleocytoviricota* viruses^5^ (a single HMM run within anvi’o). The interface considers the sequence composition, differential coverage, GC-content, and taxonomic signal of each contig, and displayed the eight hallmark genes as individual layers as well 149 additional orthologous groups often found in reference *Nucleocytoviricota* viruses^5^ as a single extra layer for guidance. During binning, no restriction was applied in term of number of giant virus core gene markers present, as long as the signal suggested the occurrence of a putative *Nucleocytoviricota* MAG. Note that while some metagenomic blocks contained a limited number of *Nucleocytoviricota* MAGs, others contained dozens. Finally, we individually refined all the *Nucleocytoviricota* and Mirus MAGs >50kbp in length as outlined in Delmont and Eren^41^, and renamed contigs they contained according to their MAG ID.

### Creation of the GOEV database

In addition to the *Nucleocytoviricota* and Mirus MAGs characterized in our study, we included marine *Nucleocytoviricota* MAGs characterized using automatic binning by Schulz et al.^7^ (n=743) and Moniruzzaman et al.^8^ (n=444), in part using *Tara* Oceans metagenomes. We also incorporated 235 reference *Nucleocytoviricota* genomes mostly characterized by means of cultivation but also cell sorting within plankton^42^. We determined the average nucleotide identity (ANI) of each pair of *Nucleocytoviricota* or Mirus MAGs using the dnadiff tool from the MUMmer package^43^ v4.0b2. MAGs were considered redundant when their ANI was >98% (minimum alignment of >25% of the smaller MAG in each comparison). Manually curated MAGs were selected to represent a group of redundant MAGs. For groups lacking manually curated MAGs, the longest MAG was selected. This analysis provided a non-redundant genomic database of 1,593 marine MAGs plus 224 reference genomes, named the GOEV database. We created a single CONTIGs database for the GOEV database using anvi’o. Prodigal^32^ was used to identify genes.

### Curation of hallmark genes

The amino-acid sequence datasets for RNApolA, RNApolB, DNApolB, and TFIIS were manually curated through BLASTp alignments (BLAST^44^ v2.10.1) and phylogenetic reconstructions, as previously described for eukaryotic hallmark genes^16^. Briefly, multiple sequences for a single hallmark gene within the same MAG were inspected based on their position in a corresponding single-protein phylogenetic tree performed with the same protocol as described above (“Diversity of DNA-dependent RNA polymerase B subunit genes” section). The genome’s multiple sequences were then aligned with BLASTp to their closest reference sequence, and to each other. In case of important overlap with >95% identity (likely corresponding to a recent duplication event), only the longest sequence was conserved; in case of clear split, the sequences were fused and accordingly labeled for further inspection. Finally, RNApolA and RNApolB sequences shorter than 200 aa were also removed, as DNApolB sequences shorter than 100 aa, and TFIIS sequences shorter than 25 aa. This step created a set of curated hallmark genes.

### Alignments, trimming, and single-protein phylogenetic analyses

For each of the four curated hallmark genes, the sequences were aligned with MAFFT^38^ v7.464 and the FFT-NS-i algorithm with default parameters. Sites with more than 50% gaps were trimmed using Goalign v0.3.5 (https://www.github.com/evolbioinfo/goalign). IQ- TREE^40^ v1.6.2 was used for the phylogenetic reconstructions, with the ModelFinder^39^ Plus option to determine the best fitting model according to BIC. Supports were computed from 1,000 replicates for the Shimodaira-Hasegawa (SH)-like approximation likelihood ratio (aLRT)^45^ and ultrafast bootstrap approximation (UFBoot^46^). As per IQ-TREE manual, supports were deemed good when SH-like aLRT >= 80% and UFBoot >= 95%. Anvi’o v7.1 was used to visualize and root the phylogenetic trees.

### Resolving hallmark genes occurring multiple times

We manually inspected all the duplicated sequences (hallmark genes detected multiple times in the same genome) that remained after the curation step, in the context of the individual phylogenetic trees (see previous section). First, duplicates were treated as putative contaminations based on major individual (i.e., not conserved within a clade) incongruences with the position of the corresponding genome in the other single-protein trees. The putative contaminants were easily identified and removed. Second, we identified hallmark gene paralogs encapsulating entire clades and/or subclades, suggesting that the duplication event occurred before the diversification of the concerned viral clades. This is notably the case for the majority of *Imitervirales*, which have two paralogs of the RNApolB. These paralogs were conserved for single-protein trees, but only the paralog clades with the shortest branch were conserved for congruence inspection and concatenation. Finally, we also detected a small clade of *Algavirales* viruses containing a homolog of TFIIS branching distantly from the ordinary TFIIS type, suggesting a gene acquisition. These sequences were not included in subsequent analyses. This step created a set of curated and duplicate-free hallmark genes.

### Supermatrix phylogenetic analysis of the GOEV database

Concatenations of the four aligned and trimmed curated and duplicated-free hallmark genes (methods as described above) were performed in order to increase the resolution of the phylogenetic tree. Genomes only containing TFIIS out of the four hallmark genes were excluded. For the remaining MAGs and reference genomes, missing sequences were replaced with gaps. Ambiguous genomes determined based on the presence of major and isolated (i.e., not a clade pattern) incongruences within single and concatenated proteins trees, as well as on frequent long branches and unstable positions in taxon sampling inferences, were removed. The concatenated phylogenetic trees were reconstructed using IQ-TREE^40^ v1.6.2 with the best fitting model according to the BIC from the ModelFinder^39^ Plus option. The resulting tree was then used as a guide tree for a phylogenetic reconstruction based on the site-specific frequency PMSF mixture model^47^ (LG+C30+F+R10). For the concatenated trees, supports were computed from 1,000 replicates for the Shimodaira-Hasegawa (SH)-like approximation likelihood ratio (aLRT)^45^ and ultrafast bootstrap approximation (UFBoot^46^). As per IQ-TREE manual, supports were deemed good when SH-like aLRT >= 80% and UFBoot >= 95%. Anvi’o v7.1 was used to visualize and root the phylogenetic trees.

### Taxonomic inference of GOEV database

We determined the taxonomy of *Nucleocytoviricota* MAGs based on the phylogenetic analysis results, using guidance from the reference genomes within the GOEV database as well as previous taxonomical inferences by Schulz et al.^7^, Moniruzzaman et al.^8^ and Aylward et al.^17^.

### Biogeography of the GOEV database

We performed a mapping of all metagenomes to calculate the mean coverage and detection of the GOEV database. Briefly, we used BWA v0.7.15 (minimum identity of 90%) and a FASTA file containing the 1,593 MAGs and 224 reference genomes to recruit short reads from all 937 metagenomes. We considered MAGs were detected in a given filter when >25% of their length was covered by reads to minimize non-specific read recruitments ^48^. The number of recruited reads below this cut-off was set to 0 before determining vertical coverage and percent of recruited reads.

### Metatranscriptomics of the GOEV database

We performed a mapping of all *Tara* Oceans metatranscriptomes to calculate the mean coverage and detection of genes found in the GOEV database. Briefly, we used BWA v0.7.15 (minimum identity of 90%) and a FASTA file containing the 0.6 million genes to recruit short reads from all 937 metagenomes.

### Viral-host predictions

Ecological network analysis was performed using a relative abundance matrix of ‘*Mirusviricota*’ MAGs in the pico-size fractions (0.22-3 μm) and relative abundances of eukaryotic genomes in the following five size fractions: 0.8-5 μm, 3-20 μm, 20-180 μm, 180-2,000 μm, and 0.8-2000 μm. To create the input files for network inference, we combined the Mirus matrix with each of the eukaryotic matrices (corresponding to different size fractions), and only the samples represented by both virus and eukaryotes were placed in new files. Relative abundances in the newly generated matrices were normalized using centered log-ratio (*clr*) transformation to Mirus and eukaryotes separately. Only MAGs observed in at least three samples were considered. Ecological network was built using FlashWeave^49^ v0.15.0 sensitive model with Julia v1.3.1. A threshold to determine the statistical significance was set to alpha = 0.01. To compare the performance of FlashWeave to naive correlations, we calculated the spearman rank correlation coefficient using the package Scipy v1.3.1. To build a global Mirus-eukaryote interactome, we pooled associations from the five size fractions by keeping the best positive or negative associations of each genome pair (i.e., the edges with the highest absolute weights). We used a phylogeny-guided filtering approach, Taxon Interaction Mapper (TIM)^18^, to predict the host using the global virus-eukaryote interactome. All the virus-eukaryote associations were mapped on the ‘*Mirusviricota*’ informational module phylogenetic tree to calculate the significance of the enrichment of specific associations using TIM. TIM provided a list of nodes in the viral tree and associated NCBI taxonomies (order, class, and phylum) of eukaryotes that show significant enrichment in the leaves under the nodes.

### Orthologous groups from Orthofinder

Orthologous groups (OGs) in Mirus MAGs (N = 111), a Mirus near-complete contiguous genome, and reference genomes from the Virus-Host Database (including 1,754 *Duplodnaviria*, 184 *Varidnaviria*, and 11 unclassified genomes) were generated. We used Orthofinder^50^ v 2.5.2 (-S diamond_ultra_sens) to generate OGs. A total of 26,045 OGs were generated and, OGs (N = 9,631) with at least five genome observations were used to cluster genomes.

### AGNOSTOS functional aggregation inference

AGNOSTOS partitioned protein coding genes from the GOEV database in groups connected by remote homologies and categorized those groups as members of the known or unknown coding sequence space based on the workflow described in Vanni et al. 2020^51^. AGNOSTOS produces groups of genes with low functional entropy as shown in Vanni et al. 2022^51^ and Delmont et al. 2022^16^ allowing us to provide functional annotation (Pfam domain architectures) for some of the gene clusters using remote homology methods.

### Identification and modeling of the Mirus major capsid protein

The putative major capsid protein of Mirus as well as the other morphogenetic module proteins were identified with the guidance of AGNOSTOS results, using HHsearch against the publicly available Pfam v35, PDB70, and UniProt/Swiss-Prot viral protein databases^52,53^. The candidate MCP was then modeled using the AlphaFold2^54,55^ and RoseTTAFold^56^. The resulting 3D models were then compared to the major capsid protein structures of phage HK97 and human cytomegalovirus and visualized using ChimeraX^57^.

### Functional inferences of *Nucleocytoviricota* genomes

Genes from the GOEV database were BLASTP against Virus-Host DB^58^, RefSeq^59^, UniRef90^60^, NCVOGs^61^, and NCBI nr database using Diamond^62^ v2.0.6 with a cut-off E-value 1 × 10^−5^. A recently published GVOG database^17^ was also used in annotation using hmmer^36^ v3.2.1 search with E-value 1 × 10^−3^ as a significant threshold. In addition, KEGG Orthology (KO) and functional categories were assigned with the Eggnog-Mapper^63^ v2.1.5. Finally, tRNAscan-SE^64^ v2.0.7 predicted 7,734 tRNAs.

### 3D structure prediction of ‘*Mirusviricota’* core gene

Proteins corresponding to ‘*Mirusviricota’* core gene clusters and lacking functional annotation based on sequence similarities were modeled using the AlphaFold2^54,55^ (-c full_dbs -t 2022-03- 12). DALI server^65^ was used to predict their functionality based on protein structure comparisons.

### Realm assignation of genes from a near complete genome

Two in-house HMM databases were created as follows. First, all coding sequences (CDS) labeled as “*Nucleocytoviricota*” were removed from the *Varidnaviria* CDS dataset (N= 53,776) in the Virus-Host Database (VHDB^66^, May 2022). To this dataset, *Tara* Ocean *Nucleocytoviricota* MAGs (all were manually curated) and 235 reference *Nucleocytoviricota* genomes were integrated. The final *Nucleocytoviricota* protein database contained 269,523 CDS. Similarly, we replaced all *Herpesvirales* CDS in the VHDB *Duplodnaviria* CDS dataset with *Herpesvirales* protein sequences downloaded from NCBI in April 2022. Additionally, a marine *Caudovirales* database including jumbo phage environmental genomes^67,68^ was integrated into the *Duplodnaviria* proteins. The final *Duplodnaviria* protein database contained 748,546 proteins. Proteins in the two databases were independently clustered at 30% sequence identity (-c 0.4 --cov-mode 5), using Linclust in MMseqs^69^ v13-45111. Gene clusters with fewer than 3 genes were removed, and the remaining gene clusters were aligned using^38^ v 7.487. HMM files (N= 16,689 and 57,259 for *Varidnaviria* and *Duplodnaviria*, respectively) were created using the hmmhuild in HMMER3^70^ v3.2.1. All proteins in the near-complete ‘*Mirusviricota*’ genome were searched against the two custom HMM databases using the hmmsearch with a cut-off E-value 1 × 10^−6^.

### Statistical analyses

A “greater” Fisher’s exact test was employed to identify KO functions as well as gene clusters with remote homologies that are differentially occurring between the 111 ‘Mirusviricota’ MAGs on one side, and all other *Nucleocytoviricota* in the GOEV database on the other side. P-values were corrected using the Benjamini-Hochberg procedure in R, and values <0.05 were considered significant.

### Naming of Mirus and Procul

The latin adjective “**Mirus**” (*surprising, strange*) was selected to describe the putative new *Duplodnaviria* phylum: the ‘***Mirusviricota’***. The latin adverb “**Procul**” (away, at distance, far off) was selected to describe the putative new class of *Nucleocytoviricota* discovered from the Arctic and Southern Oceans: the ‘***Proculviricetes’***.

## Data availability

Data our study generated has been made publicly available at https://doi.org/10.6084/m9.figshare.20284713. This link provides access to (1) the RNApolB genes reconstructed from the *Tara* Oceans assemblies (along with references), (2) individual FASTA files for the 1,593 non-redundant marine *Nucleocytoviricota* and mirusvirus MAGs (including the 697 manually curated MAGs from our survey) and 224 reference *Nucleocytoviricota* genomes contained in the GOEV database, (3) the GOEV anvi’o CONTIGS database, (4) genes and proteins found in the GOEV database, (5) manually curated hallmark genes and corresponding phylogenies, (6) HMMs for hallmark genes, (7) a FASTA file for the near-complete contiguous genome (SAMEA2619782_METAG_scaffold_2), (8) and supplemental tables.

## Contributions

Tom O. Delmont conducted the study, which was initiated alongside Morgan Gaïa and Patrick Forterre. Morgan Gaïa, Lingjie Meng, Mart Krupovic, Chiara Vanni, Eric Pelletier and Tom O. Delmont performed the primary data analysis. Tom O. Delmont completed the genome-resolved metagenomic analysis. Morgan Gaïa and Tom O. Delmont curated the marker genes and identified the biological duplicates. Morgan Gaïa performed phylogenetic and phylogenomic analyses. Lingjie Meng performed functional analyses, protein and virus-host predictions with the supervision of Hiroyuki Ogata. Chiara Vanni produced gene cluster with remote homologies with the supervision of Antonio Fernandez-Guerra. Mart Krupovic discovered the major capsid protein of ‘*Mirusviricota*’ and other key genes of the virion module. Eric Pelletier performed comparative genomic, biogeographic and metatranscriptomic analyses. All the authors contributed to interpreting the data and writing the manuscript.

## Acknowledgments

Our survey was made possible by two scientific endeavors: the sampling and sequencing efforts by the *Tara* Oceans Project, and the bioinformatics and visualization capabilities afforded by anvi’o. We are indebted to all who contributed to these efforts, as well as other open-source bioinformatics tools for their commitment to transparency and openness. *Tara* Oceans (which includes the *Tara* Oceans and *Tara* Oceans Polar Circle expeditions) would not exist without the leadership of the *Tara* Oceans Foundation and the continuous support of 23 institutes (https://oceans.taraexpeditions.org/). Some of the computations were performed using the platine, titane and curie HPC machine provided through GENCI grants (t2011076389, t2012076389, t2013036389, t2014036389, t2015036389 and t2016036389). This study was in part supported by Japan Society for the Promotion of Science (JSPS) KAKENHI (18H02279, 22H00384), Research Unit for Development of Global Sustainability, Kyoto University Research Coordination Alliance, and the International Collaborative Research Program of the Institute for Chemical Research, Kyoto University (2022-26, 2021-29, 2020-28). M.K. was supported by grants from the l’Agence Nationale de la Recherche (ANR-20-CE20-0009-02 and ANR-21-CE11- 0001-01). Part of computational work was performed at the SuperComputer System, Institute for Chemical Research, Kyoto University. This article is contribution number XXX of *Tara* Oceans.

## Supplemental figures

**Figure S1:**
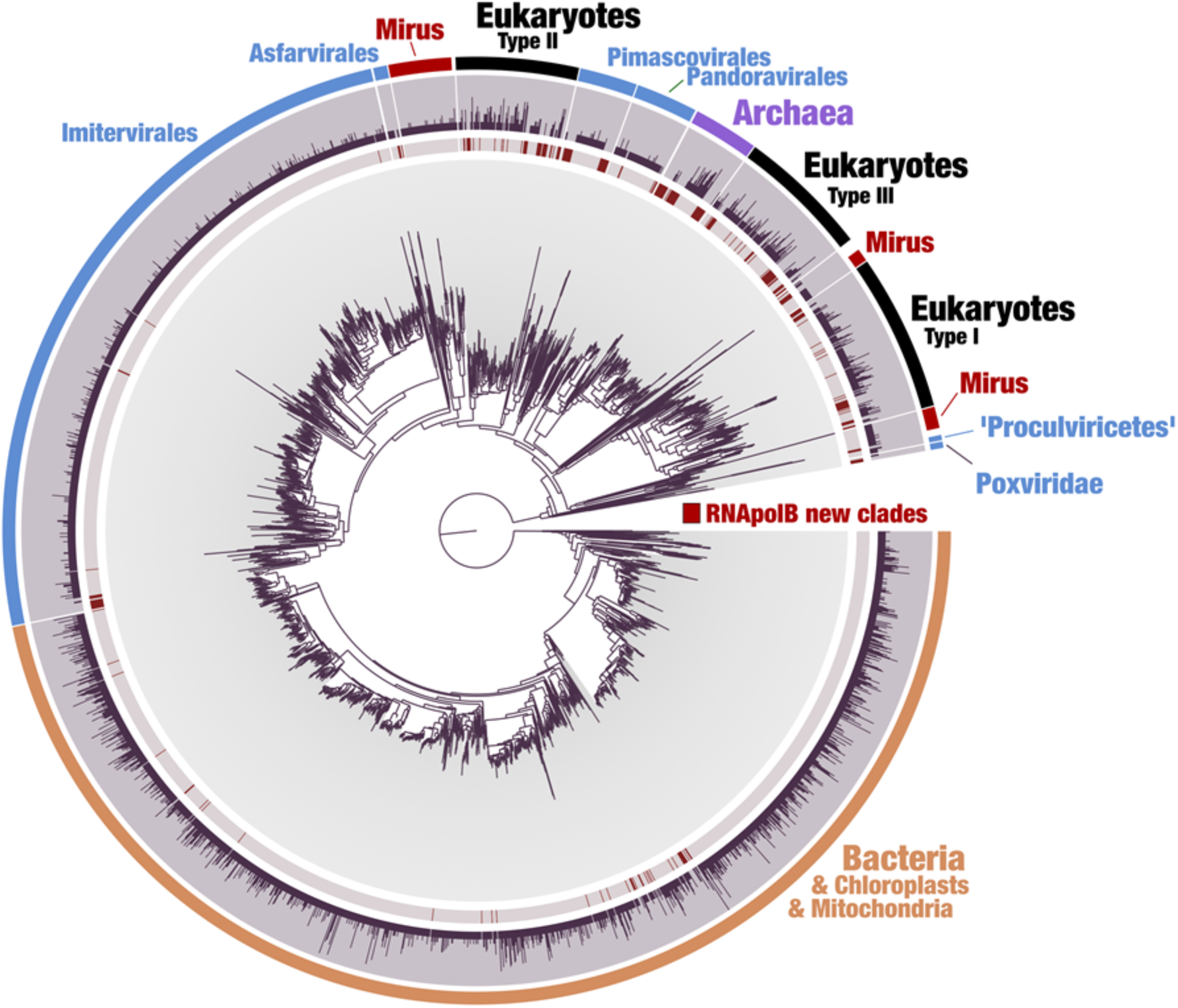
Evolutionary diversity of the DNA-dependent RNA polymerase B in the sunlit ocean. The maximum-likelihood phylogenetic tree is based on 2,728 RNApolB sequences more than 800 amino acids in length with similarity <90% (gray color) identified from 11 large marine metagenomic co-assemblies. This analysis also includes 262 reference RNApolB sequences (red color in the first layer) corresponding to known archaeal, bacterial, eukaryotic and giant virus lineages for perspective. The second layer shows the number of RNApolB sequences from the 11 metagenomic co-assemblies that match to the selected amino acid sequence with identity >90%. Finally, RNApolB new lineages are displayed in red, labelled as “Mirus”.

**Figure S2:**
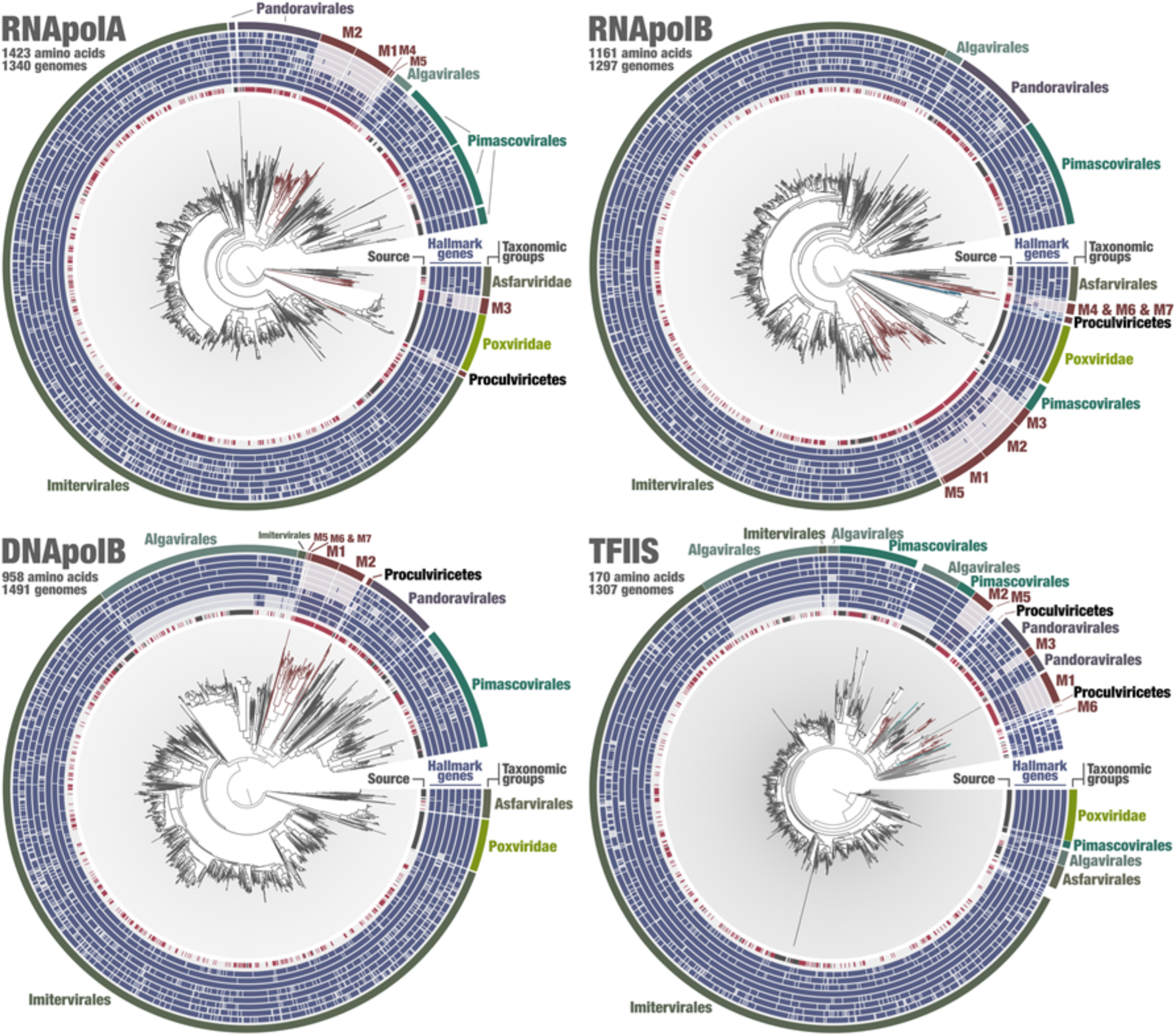
Single-protein phylogenies of the four informational hallmark genes in the GOEV database. Maximum-likelihood phylogenetic trees of the RNpolA, RNApolb, DNApolB and TFIIS. In each tree, branches corresponding to mirusviruses are colored in red. The layers provide additional information, from innermost to outermost: the source (red for the MAGs in our survey, black for references, uncolored for MAGs from other surveys), and the presence/absence of *Nucleocytoviricota* hallmark genes (RNApolA, RNpolB, DNApolB, TFIIS, MCP, primase, VLTF3, pATPase) in the corresponding genomes.

**Figure S3:**
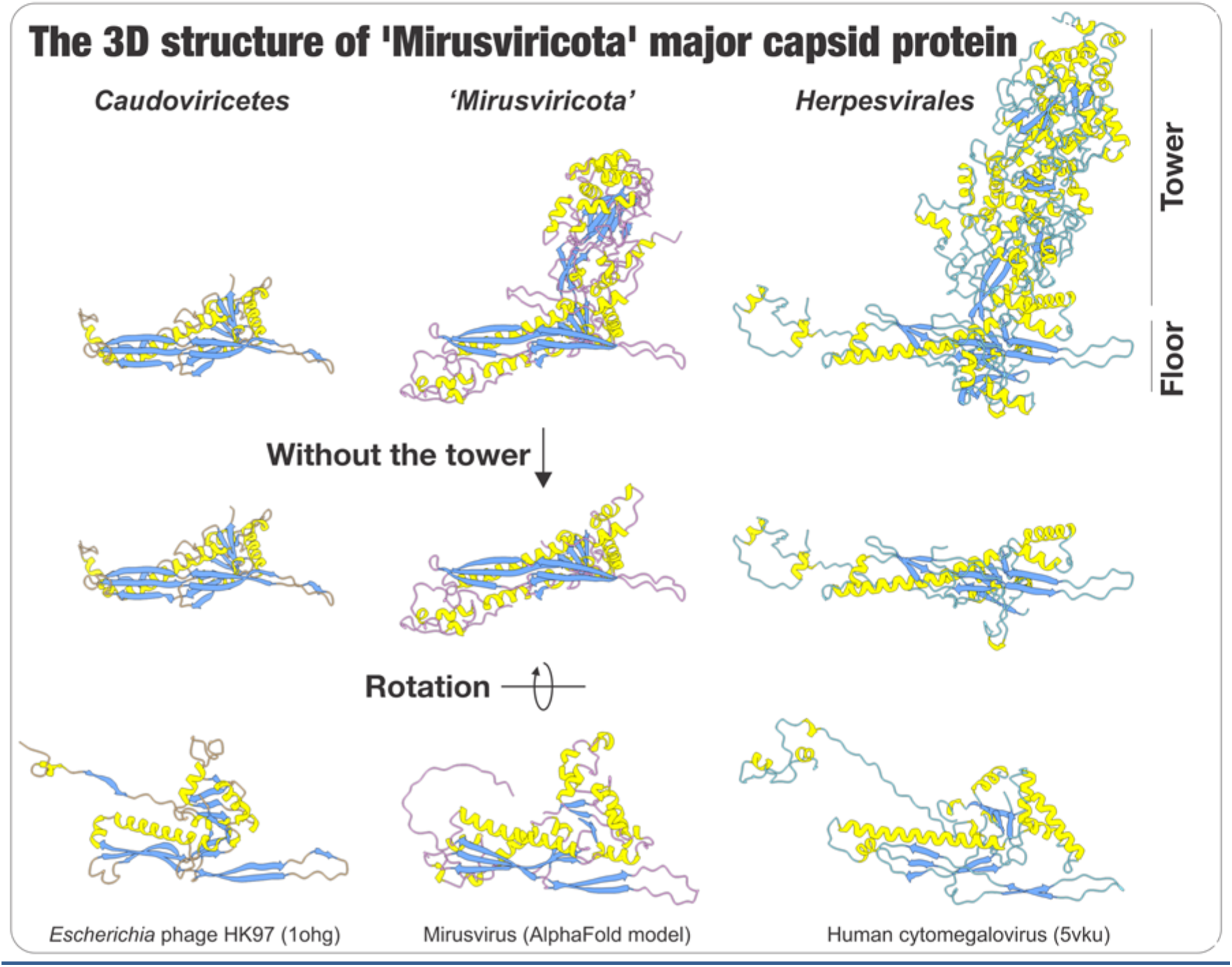
3D structure of the major capsid protein (MCP). The figure displays MCP 3D structures for *Escherichia* phage HK97 (*Caudoviricetes*), a reference genome for the mirusviruses (estimated using Alphafold), and the human cytomegalovirus (*Herpesvirales*).

**Figure S4:**
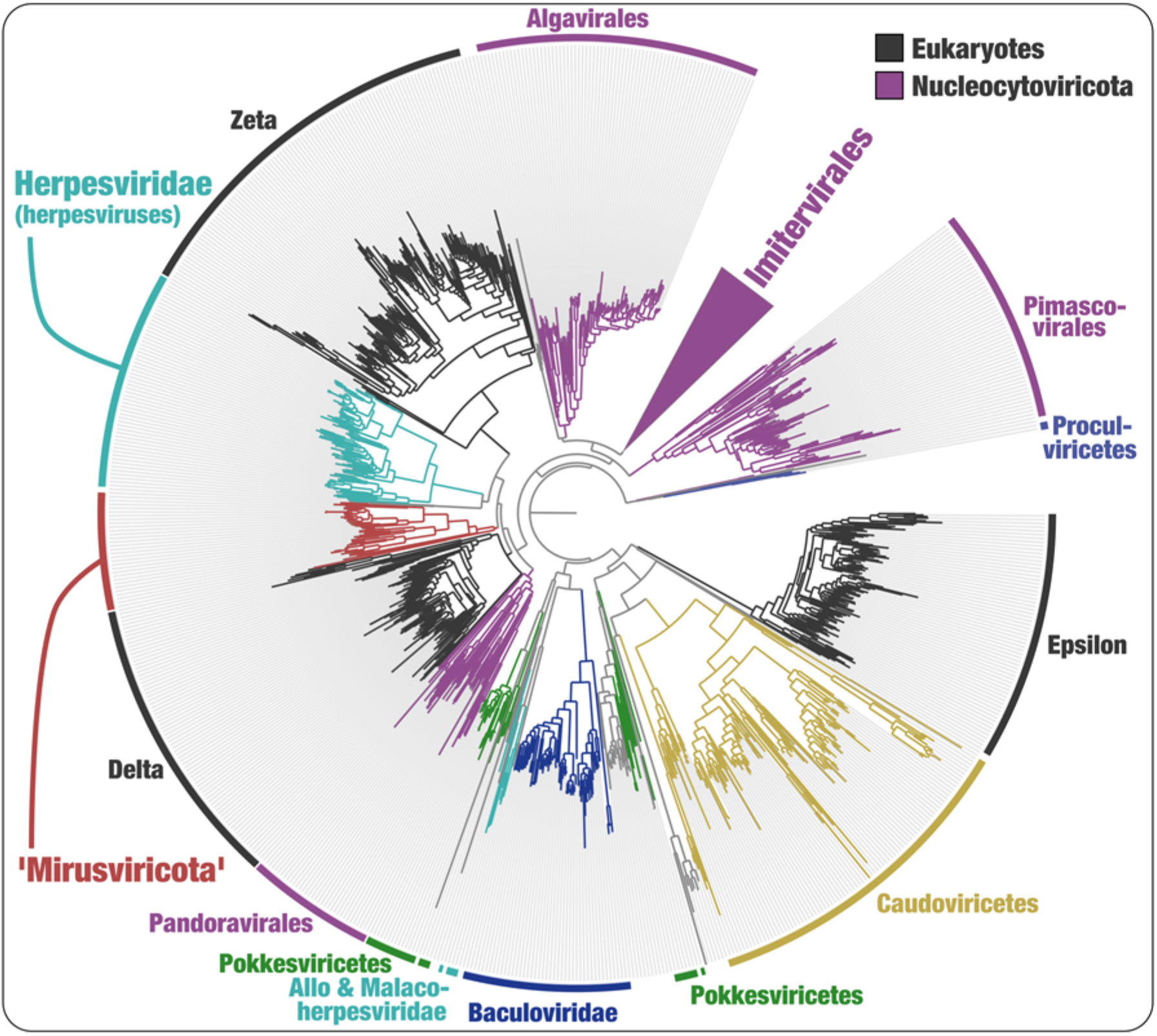
Phylogeny of the DNApolB hallmark genes. The figure displays a maximum-likelihood phylogenetic tree of DNA-polymerase B-family sequences (1,080 sites, 2,213 sequences) from the database described herein, *Duplodnaviria* and *Baculoviridae* sequences from the NCBI viral genomic database, and eukaryotic and viral sequences from Kazlauskas et al.^24^. Eukaryotic Epsilon-type and related sequences were used as outgroup.

**Figure S5:**
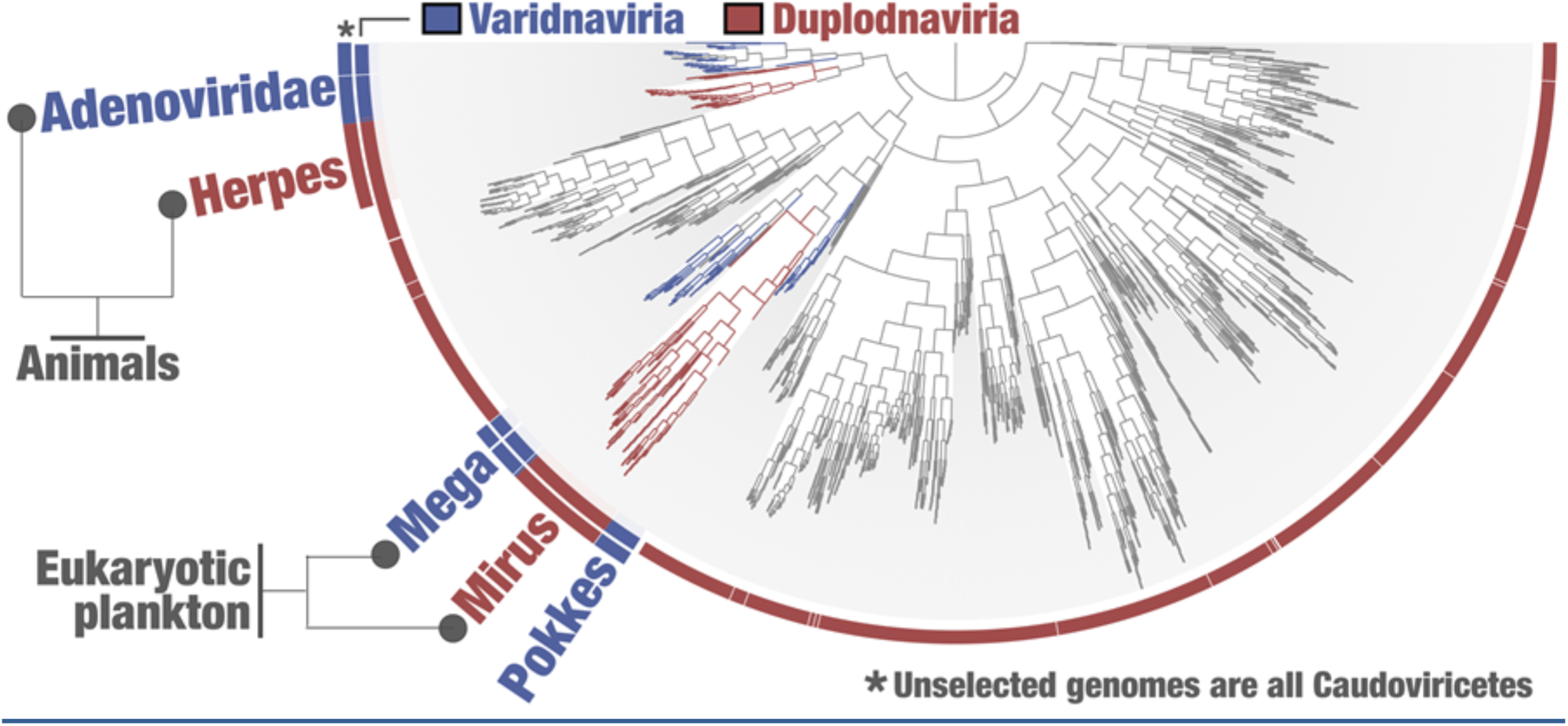
Functional clustering of mirusviruses and reference viral genomes from culture. The inner tree is a clustering of ‘*Mirusviricota*’ and other genomes based on the occurrence of all gene clusters (OrthoFinder method, Bray-Curtis distance).

**Figure S6:**
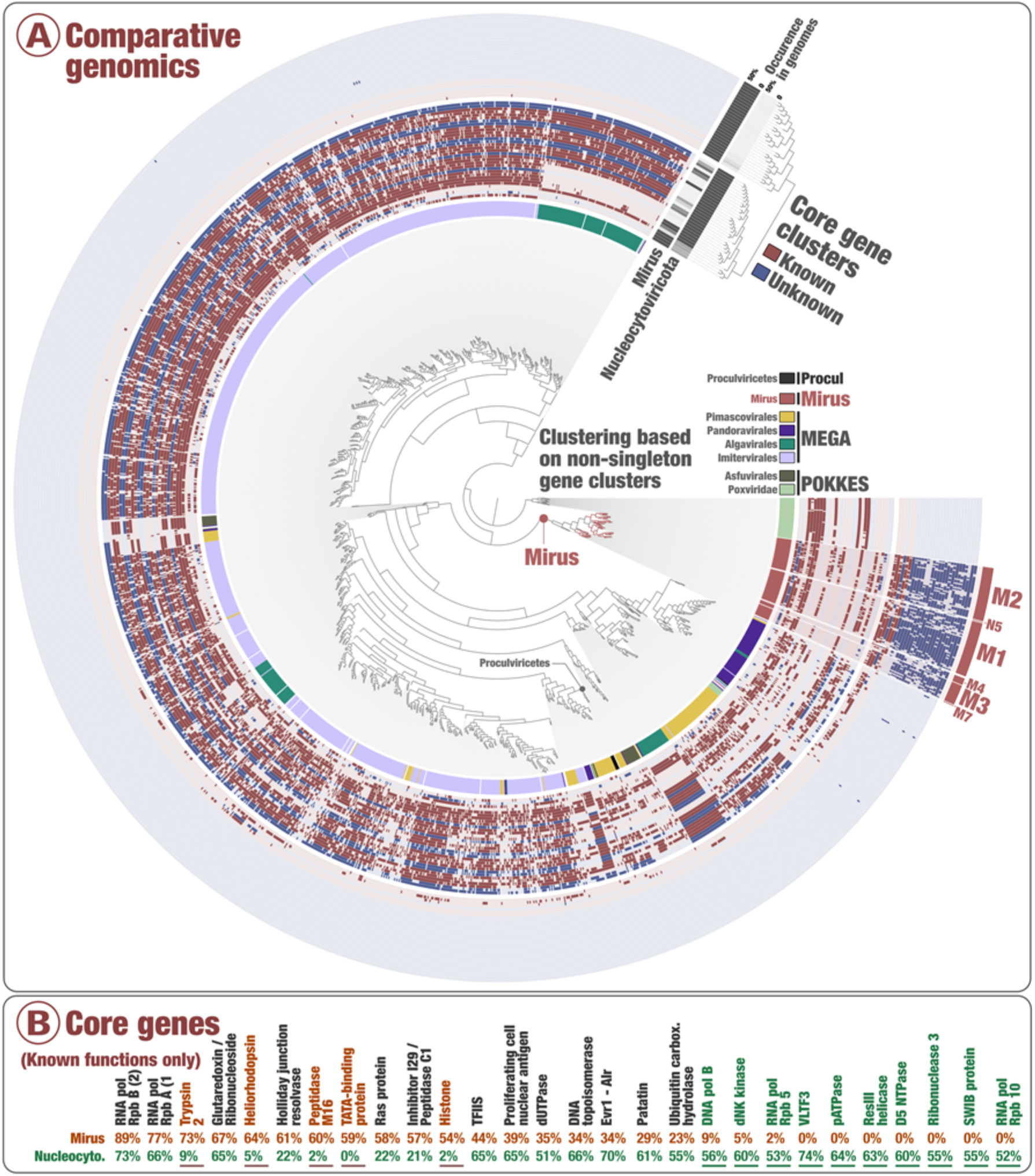
Functional clustering of abundant and widespread marine viruses within mirusviruses and *Nucleocytoviricota*. In panel A, the inner tree is a clustering of ‘Mirusviricota’ and *Nucleocytoviricota* genomes >100kbp in length based on the occurrence of all the non-singleton gene clusters (Euclidean distance), rooted with the *Chordopoxvirinae* subfamily of *Poxviridae* genomes. Layers of information display the main taxonomy of *Nucleocytoviricota* as well as the occurrence of 60 gene clusters detected in at least 50% of ‘*Mirusviricota*’ or *Nucleocytoviricota*. The 60 gene clusters are clustered based on their occurrence (absence/presence) across the genomes. Panel B displays the occurrence of gene clusters of known Pfam functions detected in at least 50% of ‘Mirusviricota’ or *Nucleocytoviricota* genomes.

**Figure S7:**
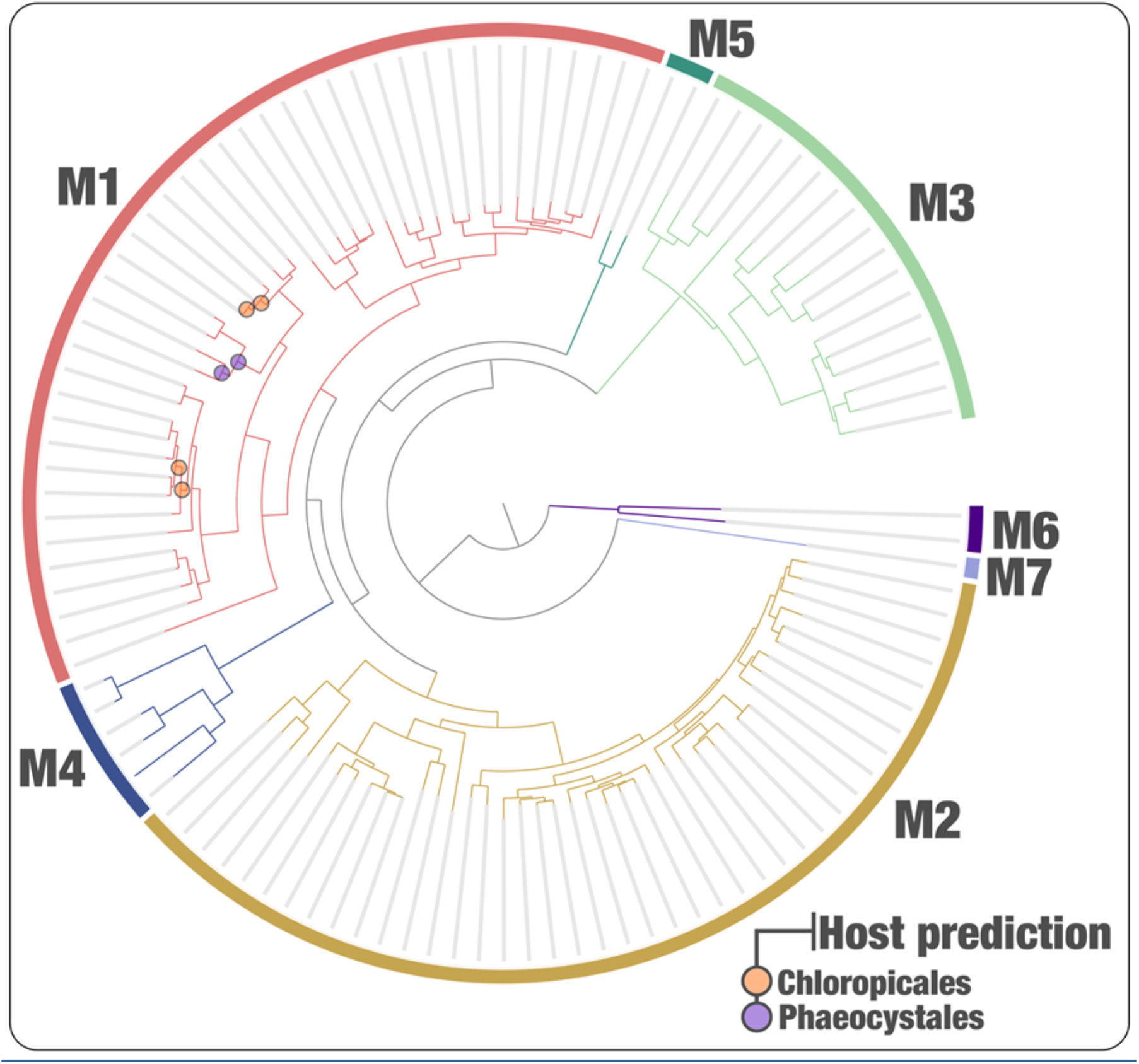
Class-level host predictions for the mirusviruses. The tree corresponds to a phylogeny of Mirus MAGs (informational module, see figure 2), and circles represents clades associated to eukaryotic lineages based on co-occurrence patterns.

**Figure S8:**
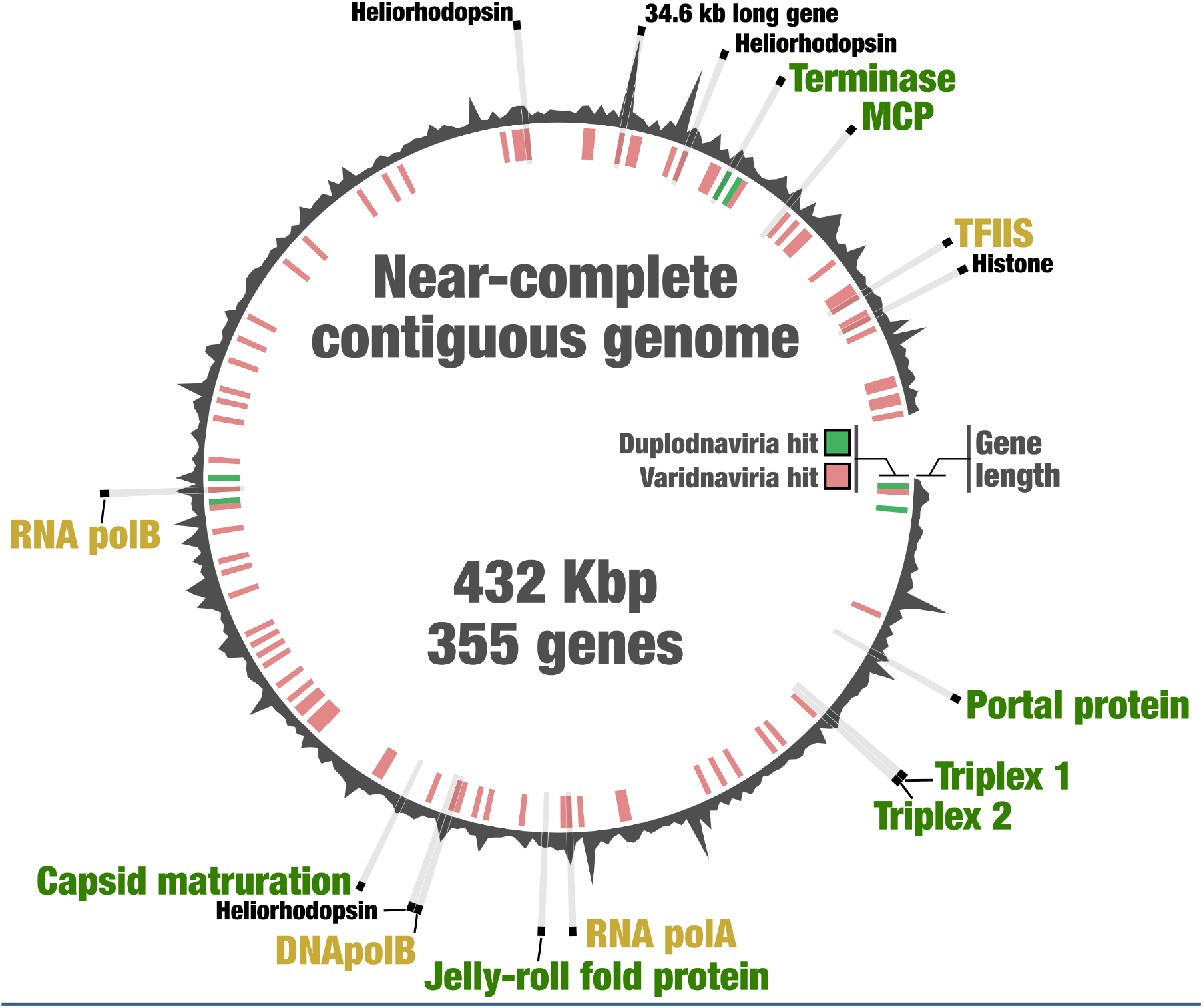
A near-complete genome for ‘*Mirusviricota’*. Syntheny of 355 genes in the mirusvirus near-complete contiguous genome highlighting the occurrence of hallmark genes for the informational and virion modules, as well as heliorhodopsins and histone. Genes with a hit to HMMs from either Duplodnaviria or Varidnaviria are labelled in green and red, respectively (inner tree).

## Supplemental tables

**Table S1:** Description of 937 Tara Oceans metagenomes.

**Table S2:** DNA-dependent RNA polymerase subunit B genes characterized from the sunlit ocean.

**Table S3:** Metabins containing DNA-dependent RNA polymerase subunit B genes of interest and used for genome-resolved metagenomics.

**Table S4:** Genomic and environmental statistics for the GOEV database.

**Table S5:** Occurrence of gene clusters with remote homologies for the GOEV database. A summary of core gene clusters and statistics are included.

**Table S6:** Occurrence of gene clusters with remote homologies for mirusviruses and reference genomes from culture that include herpesviruses. Genes shared between mirusviruses and herpesviruses are summarized.

**Table S7:** Functional annotations for genes found in the GOEV database. Statistics are included.

**Table S8:** Biogeographic signal for the GOEV database. The table includes metagenomic metadata and the mean coverage of all genomes across all metatranscriptomes,

**Table S9:** *In situ* expression of genes in the GOEV database based on 1,140 Tara Oceans metatranscriptomes. The table includes metatranscriptomics metadata, mean coverage of all Mirus genes across all metatranscriptomes, as well as individual gene and core gene expression levels when considering the ten most abundant mirusviruses are further described.

## References

1. Koonin, E. v. et al. Global Organization and Proposed Megataxonomy of the Virus World. Microbiol Mol Biol Rev 84, (2020).

2. Krupovic, M., Dolja, V. v. & Koonin, E. v. The LUCA and its complex virome. Nature Reviews Microbiology 2020 18:11 18, 661–670 (2020).

3. Koonin, E. v., Dolja, V. v. & Krupovic, M. Origins and evolution of viruses of eukaryotes: The ultimate modularity. Virology 479–480, 2–25 (2015).

4. Krupovic, M. & Koonin, E. v. Polintons: a hotbed of eukaryotic virus, transposon and plasmid evolution. Nat Rev Microbiol 13, 105–115 (2015).

5. Guglielmini, J., Woo, A. C., Krupovic, M., Forterre, P. & Gaia, M. Diversification of giant and large eukaryotic dsDNA viruses predated the origin of modern eukaryotes. Proc Natl Acad Sci U S A 116, 19585–19592 (2019).

6. Woo, A. C., Gaia, M., Guglielmini, J., da Cunha, V. & Forterre, P. Phylogeny of the Varidnaviria Morphogenesis Module: Congruence and Incongruence With the Tree of Life and Viral Taxonomy. Front Microbiol 12, 1708 (2021).

7. Schulz, F. et al. Giant virus diversity and host interactions through global metagenomics. Nature (2020) doi:10.1038/s41586-020-1957-x.

8. Moniruzzaman, M., Martinez-Gutierrez, C. A., Weinheimer, A. R. & Aylward, F. O. Dynamic genome evolution and complex virocell metabolism of globally-distributed giant viruses. Nature Communications 2020 11:1 11, 1–11 (2020).

9. Endo, H. et al. Biogeography of marine giant viruses reveals their interplay with eukaryotes and ecological functions. Nature Ecology & Evolution 2020 4:12 4, 1639–1649 (2020).

10. Mann, N. H. Phages of the marine cyanobacterial picophytoplankton. FEMS Microbiol Rev 27, 17–34 (2003).

11. Kaneko, H. et al. Eukaryotic virus composition can predict the efficiency of carbon export in the global ocean. iScience 24, 102002 (2021).

12. Gregory, A. C. et al. Marine DNA Viral Macro- and Microdiversity from Pole to Pole. Cell 177, 1109-1123.e14 (2019).

13. Laber, C. P. et al. Coccolithovirus facilitation of carbon export in the North Atlantic. Nature Microbiology 2018 3:5 3, 537–547 (2018).

14. Sunagawa, S. et al. Tara Oceans: towards global ocean ecosystems biology. Nat Rev Microbiol 1–18 (2020) doi:10.1038/s41579-020-0364-5.

15. Delmont, T. O. et al. Heterotrophic bacterial diazotrophs are more abundant than their cyanobacterial counterparts in metagenomes covering most of the sunlit ocean. The ISME Journal 2021 1–10 (2021) doi:10.1038/s41396-021-01135-1.

16. Delmont, T. O. et al. Functional repertoire convergence of distantly related eukaryotic plankton lineages abundant in the sunlit ocean. Cell Genomics 100123 (2022) doi:10.1016/J.XGEN.2022.100123.

17. Aylward, F. O., Moniruzzaman, M., Ha, A. D. & Koonin, E. v. A phylogenomic framework for charting the diversity and evolution of giant viruses. PLoS Biol 19, e3001430 (2021).

18. de Vargas, C. et al. Eukaryotic plankton diversity in the sunlit ocean. Science (1979) 348, (2015).

19. Carradec, Q. et al. A global ocean atlas of eukaryotic genes. Nature Communications 2018 9:1 9, 1–13 (2018).

20. Mihara, T. et al. Taxon Richness of ‘Megaviridae’ Exceeds those of Bacteria and Archaea in the Ocean. Microbes Environ 33, 162–171 (2018).

21. Zhang, Y. et al. Atomic structure of the human herpesvirus 6B capsid and capsid-associated tegument complexes. Nat Commun 10, (2019).

22. Duda, R. L. & Teschke, C. M. The amazing HK97 fold: versatile results of modest differences. Curr Opin Virol 36, 9–16 (2019).

23. Hua, J. et al. Capsids and Genomes of Jumbo-Sized Bacteriophages Reveal the Evolutionary Reach of the HK97 Fold. mBio 8, (2017).

24. Kazlauskas, D., Krupovic, M., Guglielmini, J., Forterre, P. & Venclovas, C. S. Diversity and evolution of B-family DNA polymerases. Nucleic Acids Res 48, 10142 (2020).

25. Paoli, L. et al. Biosynthetic potential of the global ocean microbiome. Nature 2022 1–8 (2022) doi:10.1038/s41586-022-04862-3.

26. Legendre, M. et al. Diversity and evolution of the emerging Pandoraviridae family. Nature Communications 2018 9:1 9, 1–12 (2018).

27. Talbert, P. B., Armache, K. J. & Henikoff, S. Viral histones: pickpocket’s prize or primordial progenitor? Epigenetics & Chromatin 2022 15:1 15, 1–20 (2022).

28. Hososhima, S. et al. Proton-transporting heliorhodopsins from marine giant viruses. Elife 11, (2022).

29. Weinheimer, A. R. & Aylward, F. O. A distinct lineage of Caudovirales that encodes a deeply branching multi-subunit RNA polymerase. Nature Communications 2020 11:1 11, 1–9 (2020).

30. Li, D., Liu, C. M., Luo, R., Sadakane, K. & Lam, T. W. MEGAHIT: An ultra-fast single-node solution for large and complex metagenomics assembly via succinct de Bruijn graph. Bioinformatics 31, 1674–1676 (2014).

31. Eren, A. M. et al. Anvi’o: an advanced analysis and visualization platform for ‘omics data. PeerJ 3, e1319 (2015).

32. Hyatt, D. et al. Prodigal: prokaryotic gene recognition and translation initiation site identification. BMC Bioinformatics 11, 119 (2010).

33. Li, H. & Durbin, R. Fast and accurate short read alignment with Burrows-Wheeler transform. Bioinformatics 25, 1754–1760 (2009).

34. Li, H. et al. The Sequence Alignment/Map format and SAMtools. Bioinformatics 25, 2078–2079 (2009).

35. Alneberg, J. et al. Binning metagenomic contigs by coverage and composition. Nat Methods 11, 1144–1146 (2014).

36. Eddy, S. R. Accelerated Profile HMM Searches. PLoS Comput Biol 7, e1002195 (2011).

37. Li, W. & Godzik, A. Cd-hit: a fast program for clustering and comparing large sets of protein or nucleotide sequences. Bioinformatics 22, 1658–1659 (2006).

38. Katoh, K. & Standley, D. M. MAFFT Multiple Sequence Alignment Software Version 7: Improvements in Performance and Usability. Mol Biol Evol 30, 772–780 (2013).

39. Kalyaanamoorthy, S., Minh, B. Q., Wong, T. K. F., Von Haeseler, A. & Jermiin, L.S. ModelFinder: fast model selection for accurate phylogenetic estimates. Nature Methods 2017 14:6 14, 587–589 (2017).

40. Nguyen, L. T., Schmidt, H. A., Von Haeseler, A. & Minh, B. Q. IQ-TREE: A Fast and Effective Stochastic Algorithm for Estimating Maximum-Likelihood Phylogenies. Mol Biol Evol 32, 268–274 (2015).

41. Delmont, T. O. & Eren, A. M. Identifying contamination with advanced visualization and analysis practices: metagenomic approaches for eukaryotic genome assemblies. PeerJ 4, e1839 (2016).

42. Needham, D. M. et al. Targeted metagenomic recovery of four divergent viruses reveals shared and distinctive characteristics of giant viruses of marine eukaryotes. Philosophical Transactions of the Royal Society B 374, (2019).

43. Delcher, A. L., Phillippy, A., Carlton, J. & Salzberg, S. L. Fast algorithms for large-scale genome alignment and comparison. Nucleic Acids Res 30, 2478–2483 (2002).

44. Altschul, S. F., Gish, W., Miller, W., Myers, E. W. & Lipman, D. J. Basic local alignment search tool. J Mol Biol 215, 403–410 (1990).

45. Guindon, S. et al. New algorithms and methods to estimate maximum-likelihood phylogenies: Assessing the performance of PhyML 3.0. Syst Biol 59, 307–321 (2010).

46. Hoang, D. T., Chernomor, O., Von Haeseler, A., Minh, B. Q. & Vinh, L. S. UFBoot2: Improving the Ultrafast Bootstrap Approximation. Mol Biol Evol 35, 518–522 (2018).

47. Wang, H. C., Minh, B. Q., Susko, E. & Roger, A. J. Modeling Site Heterogeneity with Posterior Mean Site Frequency Profiles Accelerates Accurate Phylogenomic Estimation. Syst Biol 67, 216–235 (2018).

48. Delmont, T. O. et al. Nitrogen-fixing populations of Planctomycetes and Proteobacteria are abundant in surface ocean metagenomes. Nature Microbiology 2018 3:7 3, 804–813 (2018).

49. Tackmann, J., Matias Rodrigues, J. F. & von Mering, C. Rapid Inference of Direct Interactions in Large-Scale Ecological Networks from Heterogeneous Microbial Sequencing Data. Cell Syst 9, 286-296.e8 (2019).

50. Emms, D. M. & Kelly, S. OrthoFinder: solving fundamental biases in whole genome comparisons dramatically improves orthogroup inference accuracy. Genome Biol 16, (2015).

51. Vanni, C. et al. Unifying the known and unknown microbial coding sequence space. Elife 11, (2022).

52. Gabler, F. et al. Protein Sequence Analysis Using the MPI Bioinformatics Toolkit. Curr Protoc Bioinformatics 72, (2020).

53. Steinegger, M. et al. HH-suite3 for fast remote homology detection and deep protein annotation. BMC Bioinformatics 20, (2019).

54. Jumper, J. et al. Highly accurate protein structure prediction with AlphaFold. Nature 2021 596:7873 596, 583–589 (2021).

55. Mirdita, M. et al. ColabFold: making protein folding accessible to all. Nat Methods 19, 679–682 (2022).

56. Baek, M. et al. Accurate prediction of protein structures and interactions using a three-track neural network. Science 373, 871–876 (2021).

57. Pettersen, E. F. et al. UCSF ChimeraX: Structure visualization for researchers, educators, and developers. Protein Sci 30, 70–82 (2021).

58. Mihara, T. et al. Linking Virus Genomes with Host Taxonomy. Viruses 8, (2016).

59. Pruitt, K. D., Tatusova, T. & Maglott, D. R. NCBI reference sequences (RefSeq): a curated non-redundant sequence database of genomes, transcripts and proteins. Nucleic Acids Res 35, (2007).

60. Suzek, B. E., Wang, Y., Huang, H., McGarvey, P. B. & Wu, C. H. UniRef clusters: a comprehensive and scalable alternative for improving sequence similarity searches. Bioinformatics 31, 926–932 (2015).

61. Yutin, N., Wolf, Y. I., Raoult, D. & Koonin, E. V. Eukaryotic large nucleocytoplasmic DNA viruses: clusters of orthologous genes and reconstruction of viral genome evolution. Virol J 6, (2009).

62. Buchfink, B., Xie, C. & Huson, D. H. Fast and sensitive protein alignment using DIAMOND. Nat Methods 12, 59–60 (2015).

63. Huerta-Cepas, J. et al. eggNOG 5.0: a hierarchical, functionally and phylogenetically annotated orthology resource based on 5090 organisms and 2502 viruses. Nucleic Acids Res 47, D309–D314 (2019).

64. Lowe, T. M. & Eddy, S. R. tRNAscan-SE: a program for improved detection of transfer RNA genes in genomic sequence. Nucleic Acids Res 25, 955–964 (1997).

65. Holm, L. & Rosenström, P. Dali server: conservation mapping in 3D. Nucleic Acids Res 38, W545 (2010).

66. Mihara, T. et al. Linking Virus Genomes with Host Taxonomy. Viruses 8, (2016).

67. Weinheimer, A. R. & Aylward, F. O. Infection strategy and biogeography distinguish cosmopolitan groups of marine jumbo bacteriophages. The ISME Journal 2022 1–11 (2022) doi:10.1038/s41396-022-01214-x.

68. Al-Shayeb, B. et al. Clades of huge phages from across Earth’s ecosystems. Nature 2020 578:7795 578, 425–431 (2020).

69. Hauser, M., Steinegger, M. & Söding, J. MMseqs software suite for fast and deep clustering and searching of large protein sequence sets. Bioinformatics 32, 1323–1330 (2016).

70. Finn, R. D., Clements, J. & Eddy, S. R. HMMER web server: interactive sequence similarity searching. Nucleic Acids Res 39, W29–W37 (2011).

